# Structure and assembly of a bacterial gasdermin pore

**DOI:** 10.1101/2023.04.20.537723

**Authors:** Alex G. Johnson, Megan L. Mayer, Stefan L. Schaefer, Nora K. McNamara-Bordewick, Gerhard Hummer, Philip J. Kranzusch

## Abstract

In response to pathogen infection, gasdermin (GSDM) proteins form membrane pores that induce a host cell death process called pyroptosis^1–3^. Studies of human and mouse GSDM pores reveal the functions and architectures of 24–33 protomers assemblies^4–9^, but the mechanism and evolutionary origin of membrane targeting and GSDM pore formation remain unknown. Here we determine a structure of a bacterial GSDM (bGSDM) pore and define a conserved mechanism of pore assembly. Engineering a panel of bGSDMs for site-specific proteolytic activation, we demonstrate that diverse bGSDMs form distinct pore sizes that range from smaller mammalian-like assemblies to exceptionally large pores containing >50 protomers. We determine a 3.3 Å cryo-EM structure of a *Vitiosangium* bGSDM in an active slinky-like oligomeric conformation and analyze bGSDM pores in a native lipid environment to create an atomic-level model of a full 52-mer bGSDM pore. Combining our structural analysis with molecular dynamics simulations and cellular assays, our results support a stepwise model of GSDM pore assembly and suggest that a covalently bound palmitoyl can leave a hydrophobic sheath and insert into the membrane before formation of the membrane-spanning β-strand regions. These results reveal the diversity of GSDM pores found in nature and explain the function of an ancient post-translational modification in enabling programmed host cell death.

## Main

In mammalian pyroptosis, active GSDMs bind the inner leaflet of the plasma membrane and assemble into pores that secrete cytokines and induce lytic cell death^1–5^. Human cells encode six GSDM proteins (GSDMA–E and PJVK), with individual GSDMs capable of targeting not only the plasma membrane but also membranes of mitochondria^10^ and intracellular bacteria^11^. Structures of human and mouse GSDM pores demonstrate that mammalian GSDM proteins form oligomeric assemblies of differing architectures with inner diameters ranging from 150–215 Å^6–9^. Recent identification of GSDM homologs encoded in invertebrates^12^, fungi^13,14^, and bacterial anti-phage defense systems^15^ further suggests that GSDMs may form a large diversity of pore structures to support distinct cellular and antiviral functions.

### bGSDM pores are large and diverse

To define the mechanism and architectural diversity of GSDM pore formation beyond closely related mammalian proteins, we reconstituted pore formation *in vitro* using a panel of bacterial GSDM (bGSDM) homologs from diverse bacterial phyla (Fig. 1 and Extended Data Fig. 1a). Human and bacterial GSDM proteins use a shared mechanism of activation where proteolytic cleavage within an interdomain linker releases a pore-forming N-terminal domain (NTD) from a partnering inhibitory C-terminal domain (CTD). Most mammalian GSDM inhibitory CTDs are ∼200 amino acids (aa) in length, but bacterial and fungal GSDMs achieve autoinhibition via a short ∼20–40 aa CTD^14,15^. GSDM activation in bacterial cells is controlled by a series of upstream proteins that sense unknown aspects of phage infection, limiting biochemical reconstitution of proteolytic activation and pore formation. We therefore sought to build on previous mammalian GSDM experiments^4,6,7,11^ to engineer bGSDMs for rational activation and enable analysis of pore formation across phylogenetically diverse systems. We first predicted bGSDM CTD cleavage sites based on our previous crystal structures of bGSDMs in inactive states^15^, and screened pairs of bGSDM full-length and ΔCTD constructs *in vivo* for toxicity in a cell growth assay (Extended Data Fig. 1a–d). For each bGSDM ΔCTD construct resulting in cellular toxicity we next purified recombinant protein variants with a series of inserted sequence-specific protease motifs in the CTD linker and used a terbium-based liposome rupture assay to identify proteins capable of full membrane pore formation *in vitro* (Fig. 1b and Extended Data Fig. 2). Using this approach, we successfully engineered controlled membrane pore formation for four bGSDM proteins from phylogenetically diverse bacteria (Fig. 1b and Extended Data Fig. 2). In agreement with previous analyses of the mammalian GSDM lipid requirements for pore formation^4,5^, we observed that human GSDMD (hGSDMD) and mouse GSDMA3 (mGSDMA3) are able to form pores in liposomes prepared from *E. coli* polar lipid extract but not phosphocholine lipids alone (Extended Data Fig. 3). In contrast, all of the engineered bGSDMs form pores in liposomes made of either simple phosphocholine lipids or *E. coli* polar lipid extract (Extended Data Fig. 3).

**Fig. 1.**
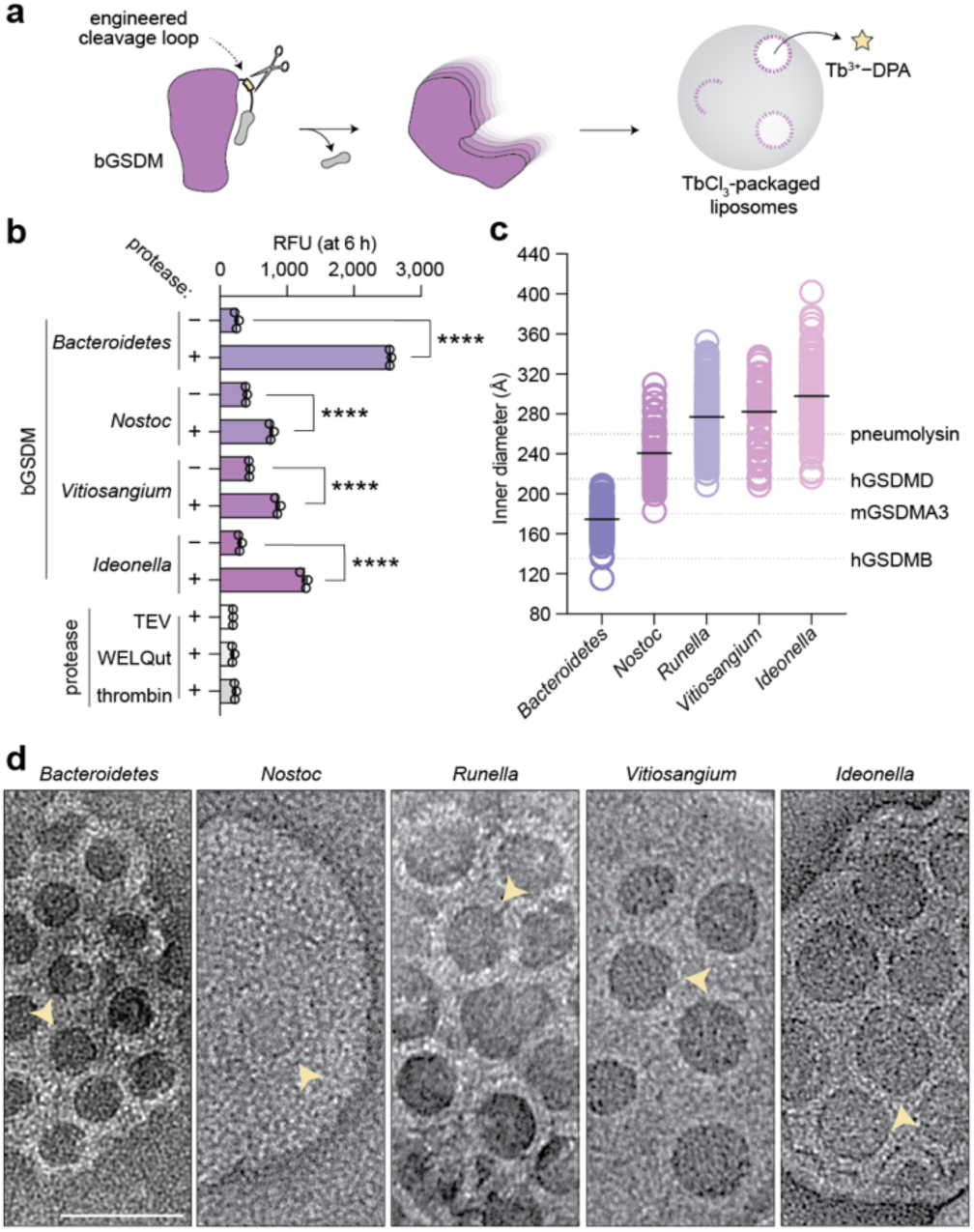
Bacterial gasdermins form pores of different sizes. **a,** Schematic of bGSDM engineering and screen for functional pore formation. Predicted cleavage site loops were mutagenized to site-specific protease cleavage motifs and tested in liposome leakage assays monitoring release of terbium chloride (TbCl_3_). **b,** Liposome leakage assay of engineered bGSDMs with matched site-specific proteases. The species of bGSDMs (and protease sites) are as follows: *Unclassified Bacteroidetes* (TEV site), *Nostoc sp. Moss4* (WELQ site), *Vitiosangium sp. GDMCC 1.1324* (thrombin), and *Ideonella sp. 201-F6* (WELQ). Error bars represent the SEM of three technical replicates. n.s. ≥ 0.05; \*\*\*\**P* < 0.0001. **c,** Size distributions of bGSDM pore inner diameters determined using negative-stain EM micrographs of pore-liposomes. The number of pores measured (n) for each species is as follows: *Bacteroidetes* (n = 117), *Nostoc* (n = 58), *Runella* (n = 189), *Vitiosangium* (n = 56), *Ideonella* (n = 123). The black bar represents the average inner diameter of measured pores. The grey dashed lines represent the average inner diameters mammalian gasdermin pores as previously measured^6,11,20^. **d,** Example negative-stain EM micrographs of bGSDM pores from five species. Scale bar = 50 nm.

We next reconstituted each successful pore-forming bGSDM variant into liposomes and visualized the membrane pores using negative-stain EM. Bacterial GSDM membrane pores are exceptionally large and diverse in architecture, spanning the range of known mammalian GSDM pores and other large pore-forming proteins (Fig. 1c,d and Extended Data Fig. 3)^16–19^. The *Bacteroidetes* bGSDM forms the smallest pores in our panel, with a mammalian-like inner diameter of ∼180 Å. In contrast, *Ideonella* bGDSM pores are massive, spanning on average ∼300 Å with some pores exceeding 400 Å and exhibiting dimensions substantially larger than the 260 Å pneumolysin cholesterol-dependent cytolysin pores produced by *Streptococcus pneumoniae* to kill target eukaryotic cells^18,19^. Notably, each bGSDM protein exhibits a distinct range of pore diameters, demonstrating that the variations in GSDM pore size are a programmed trait controlled by protein sequence specificity. Together, these results demonstrate that the size range of bGSDM pores extends substantially beyond that of characterized human GSDMs and that pore architecture is a diverse feature across GSDM protein evolution.

### Cryo-EM structural analysis of bGSDM oligomerization

To further compare the molecular basis of bacterial and human GSDM pore formation we extracted bGSDM assemblies from pore-liposome samples and performed single-particle cryo-EM analysis. We focused on *Bacteroidetes* bGSDM pores that form the smallest assemblies observed in our panel and *Vitiosangium* bGSDM pores that uniformly form some of the largest observed bGSDM assemblies. Cryo-EM and 2D classification analysis of extracted *Bacteroidetes* bGSDM pores reveals a rigid 30–31 protomer assembly that is remarkably similar to the hGSDMD 33-mer pore^20^ (Extended Data Fig. 4a,b). We confirmed assembly of predominantly 31-mer *Bacteroidetes* bGSDM pores directly in the native lipid environment, demonstrating that some bGSDM proteins form pores sharing the overall size of GSDM proteins in human immunity (Extended Data Fig. 4a,b). In contrast to the smaller *Bacteroidetes* bGSDM pores, cryo-EM analysis of *Vitiosangium* bGSDM samples demonstrated giant assemblies form through incorporation of ∼48–54 protomers (Fig. 2a and Extended Data Fig. 5a).

**Fig. 2.**
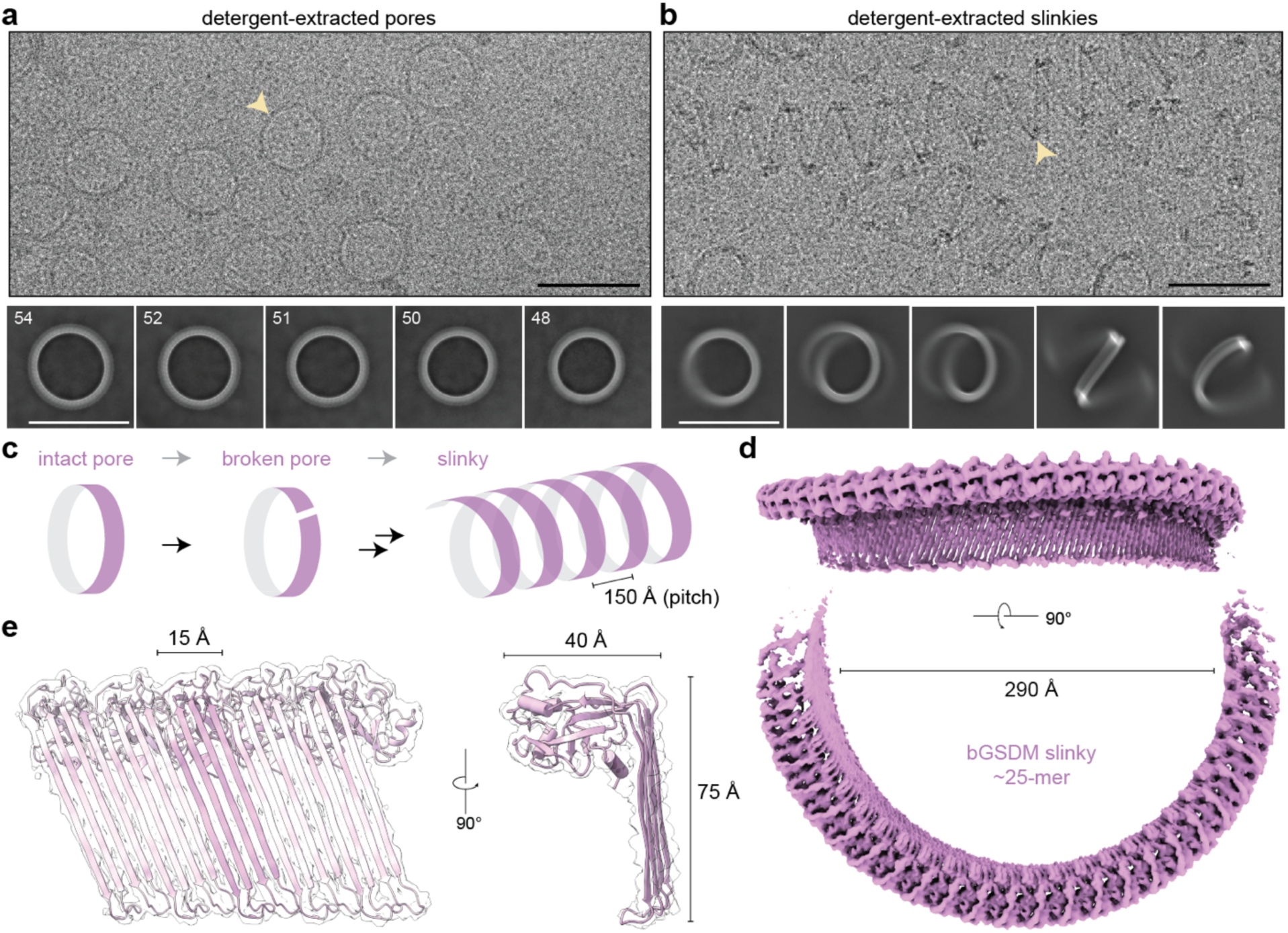
Structure of an active bGSDM oligomer from detergent extracted pores. **a,** Cryo-EM micrograph and representative 2D class averages of HECAMEG detergent-extracted *Vitiosangium* bGSDM pores. The yellow arrowhead points to an example pore. Numbers in the upper left-hand corner of 2D classes represent the number of bGSDM protomers observed in that class. Scale bars = 50 nm. **b,** Cryo-EM micrograph and representative 2D class averages of DDMAB detergent-extracted *Vitiosangium* bGSDM slinkies. The yellow arrowhead points to an example slinky. Scale bars = 50 nm. **c,** Model of structural relationship between bGSDM pores and slinky oligomers, with the pitch of a helical turn indicated. **d,** Cryo-EM map of an ∼25 protomer turn of the *Vitiosangium* bGSDM slinky. For the bottom map, the dimension of the inner diameter is shown. **e,** bGSDM active model overlaid on a portion of the slinky map. Left, 5-protomer model fit into the bGSDM slinky map. A single protomer is shown with darker magenta in the center. Right, view along the oligomerization interface of the slinky onto a single protomer. The dimensions for protomer size are shown.

Previously, X-ray crystal structural analysis of the *Vitiosangium* bGSDM revealed the unexpected self-palmitoylation of an N-terminal cysteine and defined how bGSDMs are held in an inhibited conformation prior to proteolytic activation^15^. To determine the mechanism of how proteolysis and palmitoylation lead to assembly of giant bGSDM pores, we focused on determining a high-resolution structure of *Vitiosangium* bGSDM in the active conformation. Due to orientation bias of the exceptionally large *Vitiosangium* bGSDM pores, single-particle 3D reconstruction was not possible for these samples. While screening a series of detergents and extraction methods we observed in addition to single ring pores that *Vitiosangium* bGSDM samples adopted double-ring assemblies and other open-ring assemblies including cracked slit-shaped oligomers as seen previously for human GSDM by cryo-EM, atomic force microscopy, and molecular dynamics (MD) simulation experiments^6,21,22^. Unexpectedly, in DDMAB detergent extractions we observed that open-ring assemblies of *Vitiosangium* bGSDM extend to form slinky-like oligomers with uniform radial dimensions closely mirroring the full pore (Fig. 2b,c and Extended Data Fig. 5b). *Vitiosangium* bGSDM slinkies are reminiscent of the helical oligomers of pneumolysin that form due to membrane-independent polymerization^23^. Additionally, mixtures of closed-ring pores and slinkies were observed in HECAMEG-extracted samples after prolonged storage, indicating that *Vitiosangium* bGSDM slinkies likely represent a stable non-native conformation derived from breaking closed-ring pores (Fig. 2c). Imaging *Vitiosangium* bGSDM slinkies by cryo-EM yielded 2D classes with excellent orientation distribution (Fig. 2b and Extended Data Fig. 6), and subsequent 3D classification and refinement of an ∼25-protomer turn enabled us to determine a complete 3.3 Å structure of *Vitiosangium* bGSDM in the active oligomeric conformation (Fig. 2d,e and Extended Data Figs. 5b,c and 7).

### Evolutionary conservation of GSDM activation

The *Vitiosangium* bGSDM active-state cryo-EM structure reveals dramatic conformational rearrangements and an ancient mechanism of activation shared between bacterial and human GSDM proteins. Using DALI^24^ to compare the *Vitiosangium* bGSDM model against all structures in the PDB, we observed active-state structures of hGSDMD and mGSDMA3 as the top hits confirming that the *Vitiosangium* slinky-derived conformation corresponds to the bGSDM active oligomeric state (Fig. 3a). Notable structures more distantly related to the *Vitiosangium* bGSDM active-state include the inactive conformation of mammalian GSDMs and members of the membrane attack complex perforin-like/cholesterol dependent cytolysin (MACPF/CDC) family of pore-forming proteins.

**Fig. 3.**
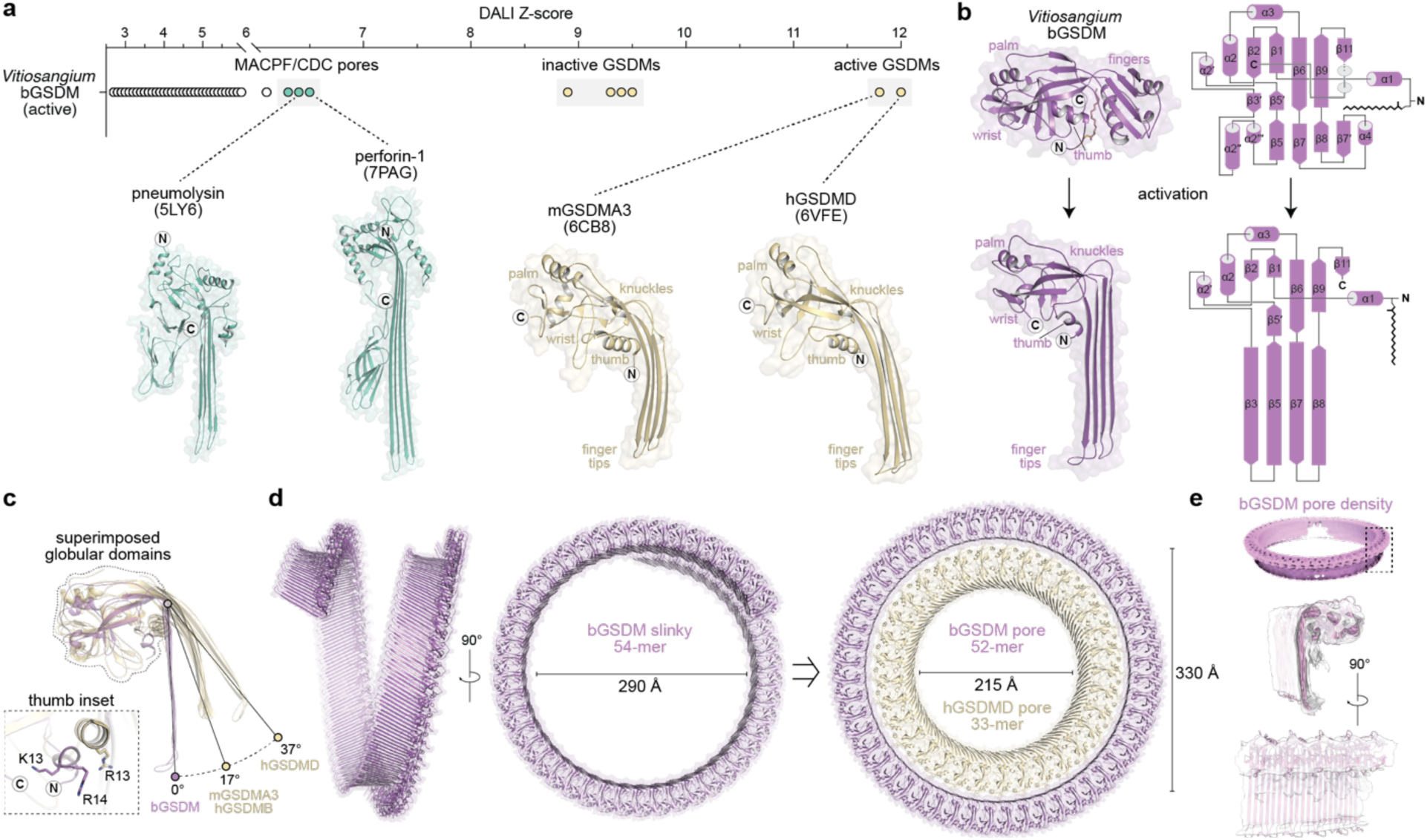
Conservation of the gasdermin active state structure from bacteria to mammals. **a,** Top, DALI Z-scores from searching the active *Vitiosangium* bGSDM structure against the full Protein Data Bank (PDB). PDB depositions with multiple similar chains were manually removed to show only unique hits. Below the graph are representative pore-forming protein structures from the membrane attack complex/perforin superfamily (MACPF) and cholesterol-dependent cytolysins (CDCs), in addition to active mammalian gasdermins. Pneumolysin and perforin structures have been reduced by 60% in size. **b,** Inactive to active state transition of the *Vitiosangium* bGSDM structure. Left, transition from the crystal structure inactive state (PDB ID 7N51) to the active cryo-EM structure (this study). Right, topology diagrams of the inactive and active structures labeling α-helices and β-sheets based on mammalian GSDM numbering^6^. The putative orientation of the palmitoyl modification is indicated in the active model. **c,** The active *Vitiosangium* bGSDM structure superimposed onto the mammalian GSDM structures from (a) and active hGSDMB (PDB ID 8ET2). Structures were superimposed by their globular domains and angles were measured using an axis at the base knuckle of the bGSDM to points at the fingertips of each structure. The inset shows the orientations of the ⍺1 thumbs in the bGSDM and the mGSDMA3 structures. Select positively charged residues are indicated, including R13 of mGSDMA3 has been proposed to interact with the cardiolipin headgroup^6^. **d,** Views of *Vitiosangium* bGSDM slinky and pore models. On the far right, the bGSDM pore is shown encompassing the hGSDMD pore model (PDB ID 6VFE). **e,** Top, a 6.5 Å resolution cryo-EM map of an elliptical closed-ring *Vitiosangium* bGSDM pore. Boxed area indicates a region where a 5-mer of the bGSDM pore model was fit into the map (below).

Strikingly, despite <15% sequence identity, the *Vitiosangium* bGDSM active-state model reveals strict conservation of the final overall ‘hand-shaped’ architecture of active mammalian GSDM proteins including the α1 ‘thumb’, the globular ‘palm’, the β1–β2 ‘wrist’ loop, and the membrane-spanning ‘fingers’ (Fig. 3b)^6,7^. Previous comparisons of the mammalian GSDM active-state structures revealed a key difference in the protein region between the globular domain and fingers region that controls the bend of the beta-sheet fingers and directs the pitch of membrane insertion^6,7^. In contrast to the gentle bend of the ‘knuckles’ regions of the hGSDMD and mGDSMA3 active-state structures, the fingers and palm of the bGSDM active-state structure intersect at a nearly perpendicular angle and form a flat plateau from the inner palm to the knuckles (Fig. 3b). Superposition of the globular domains of the bGSDM and mammalian GSDM structures demonstrates that bGSDM forms the sharpest angle with respect to the palm region, rotating 37° inward compared to the hGSDMD conformation (Fig. 3c). This rotation compresses the bGSDM α1 thumb into the globular domain, where the N- and C-termini are held in unusually close proximity (Fig. 3c). Unlike the mammalian GSDM fingers whose β-strands differ in length from 15–22 aa, the bGSDM fingers are nearly uniform at 17–18 aa each, and the fingertips, which resemble 2–7 aa β-hairpins in mammalian GSDMs, are expanded into 8–11 aa loops (Fig. 3b,c). Importantly, multiple other bGSDM sequences, including the *Lysobacter enzymogenes* bGSDM involved in anti-phage defense, are each readily modeled using the *Vitiosangium* bGSDM active-state structure supporting that the conformational changes controlling bGSDM activation are likely conserved across diverse phyla (Extended Data Fig. 8a).

Comparison of the *Vitiosangium* bGSDM inactive-state crystal structure with the active-state cryo-EM model additionally explains bacteria-specific transitions that direct activation. Whereas the mammalian GSDM fingers form almost exclusively through the elongation of existing β-strands, the bGSDM fingers form via the local refolding of several alpha-helices (α2′′, α2′′′, and α4) and the β7′ strand, which are largely absent in mammalian GSDM structures (Fig. 3b and Extended Data Fig. 8b,c). In particular, the α2′′ helix contributes multiple residues to the hydrophobic groove housing the palmitoylated N-terminal cysteine residue that stabilizes the bGSDM autoinhibited state. Though no clear electron density could be observed for the cysteine in the active structure, reorientation of the α1 thumb is predicted to flip the covalently bound palmitoyl for insertion into the surrounding lipid membrane (Fig. 3b).

Like members of the MACPF/CDC family^16–18^, GSDMs assemble into large membrane-spanning β-barrels wherein each protomer contributes four β strands to the complete pore. Despite this similarity, a previous analysis concluded that GSDMs were unlikely to have arisen by divergent evolution from the MACPF/CDC proteins, in part, due to differences in pore-forming mechanisms and the absence of conserved glycines at the knuckle region of the GSDM structures^6^. We confirmed that these glycines are largely absent from the bGSDMs and also did not detect notable homology in the structures of the bGSDM and pneumolysin active state models outside of the central β strand region (Extended Data Fig. 8b,c and Extended Data Fig. 9a). However, the above noted bGSDM-specific refolding of domains which form the central β3 and β5 strands suggests that substiantial variations in the mechanism of pore-formation of distant-related GSDMs are possible. We therefore hypothesized that features or shared homology between GSDMs and MACPF/CDC proteins might be hidden in more distantly-related bGSDM homologs. To test this hypothesis, we used FoldSeek^25^ to query the AlphaFold database and uncovered a surprising cluster of bGSDM-like proteins with cytolysin-like features (Extended Data Fig. 9b,c). These proteins are encoded by multiple *Gammaproteobacteria* including *Pseudomonas* and *Lysobacter* species (>200 homologs on the IMG database) and have predicted homology to the pore-forming domain of bGSDMs but not MACPF/CDC proteins. Intriguingly, unlike the bGSDMs studied so far, these proteins are not encoded by operons with proteases and are C-terminally fused to immunoglobulin(Ig)-like β-sandwich domains with homology to the cholesterol recognition domains of pneumolysin and other cytolysins^19^. The direct fusion of the bGSDM-like pore-forming domain with an Ig-like β-sandwich domain raises the possibility that bGSDMs share a common ancestor with a cytolysin-like pore-forming toxin. We therefore propose that GSDM self-intoxication arose by divergent evolution from ancient pore-forming toxins that target non-self membranes.

To visualize the *Vitiosangium* bGSDM in its natural closed-pore state, we assembled a 54-mer structure based on the radius of the slinky oligomeric conformation and then realigned protomers into a circular 52-mer geometric model corresponding to the dominant size observed in 2D classification of intact pores (Fig. 3d). The resulting model supports continuous hydrogen bonding around a giant pore formed by 208 β-strands. Notably, the *Vitiosangium* bGSDM pore has an inner diameter of 330 Å that is wide enough to completely encompass the largest known mammalian pore of hGSDMD (Fig. 3d). We further demonstrated that most residue-residue contacts at the subunit interface are preserved between the slinky-like oligomer and the circular pore (Extended Data Fig. 11a). To verify that the model represents the conformation of the closed-ring pore, we optimized the extraction and vitrification of *Vitiosangium* bGSDM pores and identified a condition wherein pores could be observed with minimal orientation bias (Extended Data Fig. 10a,b). A dataset collected on this sample revealed 3D classes of closed-ring structures of multiple sizes, a double-ring pore, and a slinky-like structure (Extended Data Fig. 10b,c). Refinement of the dominant class yielded a map of the flexible pore assembly at 6.5 Å resolution that is slightly oval in shape (Extended Data Fig. 10c). We docked an elliptical 52-mer of the slinky-derived pore structure and confirmed that all major features of the the model closely match the oligomeric conformation of the closed-ring pore (Fig. 3e and Extended Data Fig. 10d,e).

We next introduced mutations to *Vitiosangium* bGSDM residues at key regions including along the *Vitiosangium* bGSDM oligomeric interface and used a cell death assay to analyze bGSDM-mediated pore formation in bacterial cells. We observed that all 17 *Vitiosangium* bGSDM single substitution mutants tested retained the ability to induce cell death, and confirmed that five of these mutants can be purified as stable proteins and rupture liposomes *in vitro* (Extended Data Fig. 11b–d and Extended Data Fig. 12a–b). Combining single mutants with the strongest effects, we identified double mutants that ablated the ability of *Vitiosangium* bGSDM to induce cell death (Extended Data Fig. 11c–d). These results indicate that bGSDM pore assembly is remarkably stable and robust to minor perturbations across the protein length— a feature distinct from mammalian GSDM counterparts^6,20^

### Mechanism of bGSDM membrane assembly

The failure of single mutations in conserved regions and at the oligomeric interface to inhibit bGSDM-mediated cell death suggests that bGSDM pore formation is a robust and energetically favorable process. Previously, X-ray crystal structures of *Vitiosangium* and other bGSDMs revealed palmitoylation of a conserved N-terminal cysteine residue that is also required for bGSDM-mediated cell death^15^. We therefore hypothesized that palmitoylation may contribute to the mechanism of membrane pore formation. To determine the function of *Vitiosangium* bGSDM palmitoylation, we mutated the conserved cysteine and residues in the surrounding pocket of the inactive structure and tested the function of these bGSDM variants for the ability to induce bacterial cell death. We observed that most mutations of the palmitoyl binding pocket did not compromise bGSDM-mediated cell death, including mutations of V89 and F111 which rearrange into the active state β3–β5 fingertip loop and knuckles of the β5 strand, respectively (Fig. 4a,b and Extended Data Fig. 11). However, we found that alanine mutations of conserved L211 and F213 residues located between the β8 and β9 strands in the active state partially disrupted bGSDM function, and a C4A mutation to the modified cysteine residue itself resulted in bGSDM inactivation and near-complete rescue of bacterial growth (Fig. 4a,b). We next purified C4A mutants of *Vitiosangium* and *Nostoc* bGSDMs and analyzed both proteins *in vitro* with liposome rupture assays. Similar to our previous analysis of the *Runella* bGSDM^15^, the *Vitiosangium* and *Nostoc* bGSDMs purified as stable proteins but each were significantly reduced in the ability to form membrane pores and cause liposome leakage (Fig. 4c). These results demonstrate that the N-terminal palmitoyl modification plays a direct role in bGSDM pore formation (Fig. 4c).

**Fig. 4.**
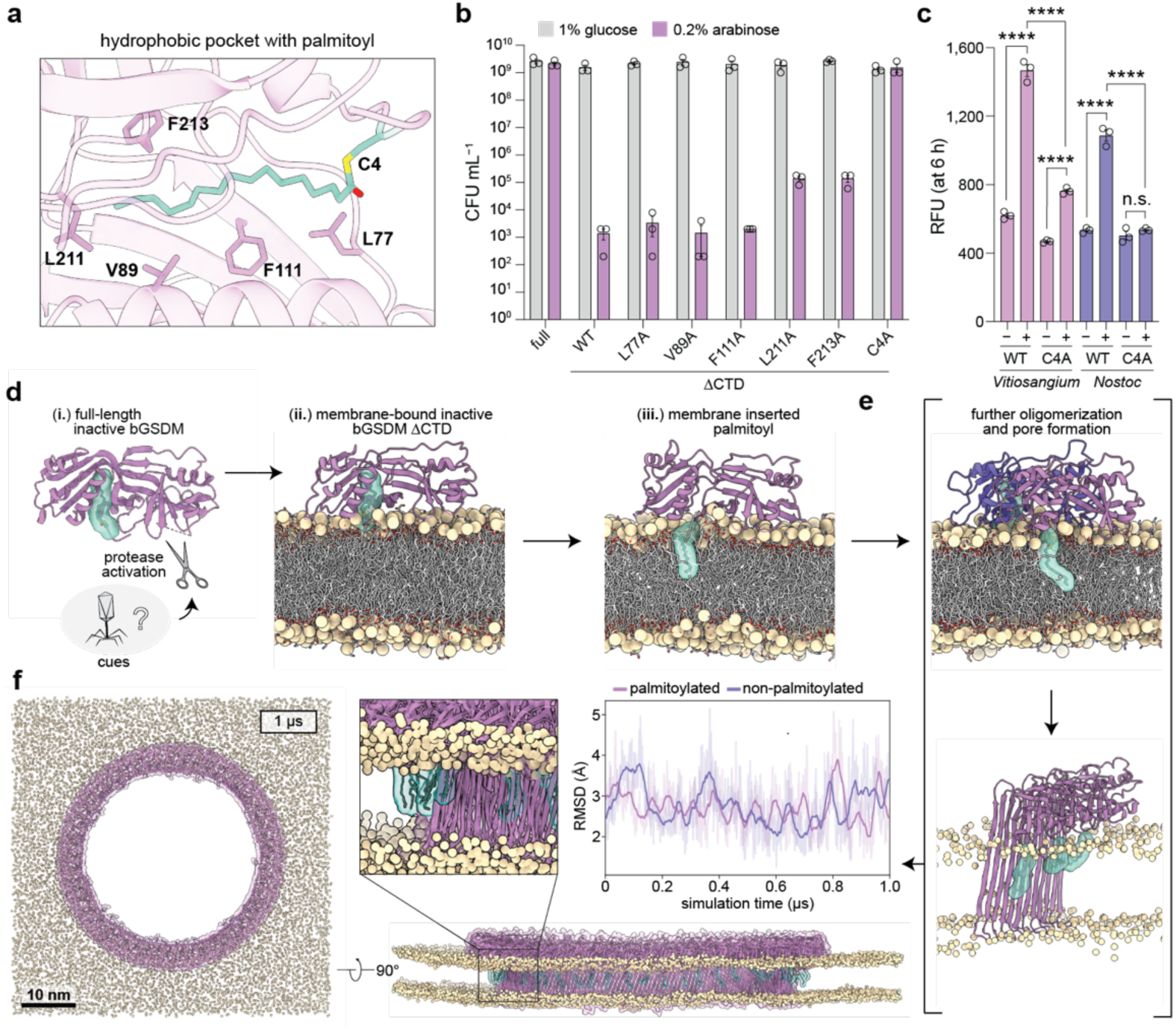
Model for membrane insertion and pore formation of palmitoylated bGSDMs. **a,** Hydrophobic pocket surrounding the N-terminal palmitoylated cysteine residue of the Vitiosangium bGSDM in the inactive state (PDB ID 7N51). Select hydrophobic residues and the palmitoylated cysteine residue (C4) are indicated. **b,** The N-terminal cysteine is critical for cell toxicity bacteria, while residues that surround it in the hydrophobic pocket are not. Colony forming units (CFUs) per mL of E. coli grown on LB-agar plates (Extended Data Fig. 9). CFU were determined from bacteria harboring plasmids encoding the full-length bGSDM (full), the N-terminal pore-forming domain alone (ΔCTD), or single mutations of the ΔCTD grown in triplicate. **c,** The N-terminal cysteine is critical for efficient pore formation in vitro. Liposome leakage assay of WT and N-terminal cysteine mutated (C4A) Vitiosangium and Nostoc bGSDMs. Engineered bGSDMs were paired with their matched site-specific protease. Error bars represent the SEM of three technical replicates and statistical significance was determined by one-way ANOVA and Tukey multiple comparison test. n.s. ≥ 0.05; ****P < 0.0001. **d,** Structural transitions during bGSDM activation. Schematic shows the anchoring process starting from (i) the inactive crystal structure of the bGSDM (PDB ID 7N51) before proteolytic cleavage, followed by snapshots showing the endpoints of MD simulations of the inactive ΔCTD adhered to the membrane (ii) at 37°C, with the palmitoyl still in the hydrophobic sheath, and (iii) at 97°C, where the palmitoyl spontaneously inserted into the membrane. **e,** Schematic of unresolved intermediate steps of oligomerization and membrane insertion represented with a simulation snapshot of a membrane-adhered dimer of ΔCTDs after 1 µs of high temperature MD simulation (top) and an oligomer in pore conformation (bottom). **f,** MD simulation of the fully assembled 52-mer pore with and without palmitoylated C4. Snapshot of the pore in top view (left) and front view (bottom right) after 1 μs of simulation. Solvent and membrane lipid tails are omitted for clarity. (top, right) RMSD fluctuations of the palmitoylated and the non-palmitoylated systems with respect to the average structure of the last 900 ns of the simulation (transparent lines) and smoothed fluctuations (opaque lines; smoothed using a 30 ns window).

We hypothesized that the palmitoyl group stimulates intermediate steps during bGSDM pore assembly. To test this hypothesis, we conducted atomistic MD simulations of the early and late steps of pore assembly (Fig. 4d–f). Starting from the full-length inactive *Vitiosangium* bGSDM crystal structure (*i*), we removed the 29 aa CTD and placed the inactive ΔCTD structure on a model bacterial membrane (*ii*). At 37°C, this structure was remarkably stable in three 2-µs long simulations, with a Cα root mean squared distance (RMSD) of ∼2 Å from the initial structure and the palmitoyl remaining sheathed in the hydrophobic binding pocket. The ΔCTD monomers and dimers remained stable also at 97°C (ΔCTD RMSD of ∼5 Å), including the proto-finger region and palmitoyl binding pocket. Nevertheless, the palmitoyl spontaneously left the hydrophobic sheath to integrate into the bulk membrane within 1 µs in 12 out of 24 monomer and 9 out of 24 dimer simulations (Fig. 4d (iii)), subsequently being replaced in some simulations by a lipid acyl chain. In simulations of a single active state protomer and of 2–3-meric bGSDMs in the membrane, the bGSDMs formed stable coiled assemblies. For the trimer, narrow membrane-spanning and water-filled channels formed, similar to those observed for hGSDMD^22^ (Fig. 4e and Extended Data Fig. 12). C4 palmitoylation did not noticeably affect the stability or shape of these small, possibly sublytic, pores. Finally, we simulated a complete 52-mer pore within a membrane. The pore was remarkably stable (RMSD of ∼3 Å) with or without palmitoylation (Fig. 4f and Supplementary Video 1). Taken together, our simulations suggest that palmitoyl membrane insertion likely precedes the structural reorganization of the bGSDM proto-finger region. Integration of the covalently attached fatty-acid tail may ensure stable membrane binding of the vulnerable, partially unfolded state and drive the system towards pore formation. Once the β-sheet is formed and membrane inserted, the palmitoyl appears to play a subordinate role. The structural similarity of the hydrophobic cavity holding the palmitoyl in different bGSDMs^26^ suggests a similar mechanism is shared in other species. To what extent the *Vitiosangium* bGSDM oligomerizes before it inserts into the membrane and whether it can form circular pre-pores, as proposed for eukaryotic GSDMs^20,27^, awaits further investigation.

Our study defines an ancient structural mechanism of GSDM pore formation. Comparison of the cryo-EM structure of the *Vitiosangium* bGSDM active-state with a previously-determined crystal structure^15^, allowed us to model how bacteria-specific structures of the inactive state transform into membrane-spanning β-sheets to form huge ∼1.3 MDa pores. Reconstitution of pore formation using five diverse bGSDMs demonstrates that pore size is heterogeneous in nature and intrinsically determined by protein sequence. Mammalian GSDM pores have well-defined roles in secretion of interleukin signals^20^. Though interleukins are not conserved in bacteria, the divergent sizes observed for bGSDM pores suggest other molecules may be released to provide an additional layer of complexity to bGSDM-mediated anti-phage defense^28^. Whereas mammalian GSDMs and pore-forming toxins recognize specific lipids in target membranes^4,5,29^, a remarkable feature of bGSDMs is the ability to form pores on membranes with simple lipid compositions^15^. This lack of specificity may reflect the absence of selective pressures on bGSDMs to distinguish the plasma membrane from the organelle membranes present in eukaryotes, or self from non-self membranes required for pore-forming toxins. Consistent with this observation, the α1 thumb, which in mGSDMA3 mediates interactions with the cardiolipin head group via R13^6^, is reoriented in the active-state bGSDM and the analogous residue of the *Vitiosangium* bGSDM is not required for GSDM-mediated cell death or liposome rupture (Fig. 3c, Extended Data Fig. 11, and Extended Data Fig. 12). This finding suggests that cardiolipin-specific interactions are not conserved features of GSDM pores, despite the prevalence of cardiolipin in bacterial membranes. Of note, we found that both bGSDMs and hGSDMD can kill bacterial cells lacking cardiolipin (Extended Data Fig. 1c)^30^. Nevertheless, future studies should investigate whether phylogenetically diverse bGSDMs are customized to the complex protein and lipid features of native bacterial membranes^31^. Besides extrinsic membrane features, bGSDMs appear to broadly utilize intrinsic properties such as palmitoylation to strengthen membrane interactions and induce membrane disorder to lower the energy barrier for pore formation. Building on the identification of bGSDM palmitoylation^15^, two recent studies describe a requirement of enzyme-dependent palmitoylation of human and mouse GSDMD for pore formation in cells^32,33^. Though the palmitoylated cysteine (C191 in humans) is located within the fingertip region instead of on the thumb in bGSDMs, we suggest that our model for palmitoyl membrane insertion may explain why the modification is conserved across the tree of life.

## Acknowledgements

The authors are grateful to members of their labs for helpful discussions, S. Rawson, S. Sterling, R. Walsh, M. Yip, and S. Shao (HMS) for advice on cryo-EM, A. Lu for assistance with protein purification, and the Max Planck Computing and Data Facility (MPCDF) for computational resources. Cryo-EM data were collected at the Harvard Cryo-EM Center for Structural Biology at Harvard Medical School and at PNCC supported by NIH grant U24GM129547. The work was funded by grants to P.J.K. from the Pew Biomedical Scholars program, the Burroughs Wellcome Fund PATH program, The Mathers Foundation, The Mark Foundation for Cancer Research, the Parker Institute for Cancer Immunotherapy, and the National Institutes of Health (1DP2GM146250-01). A.G.J. is supported through a Life Science Research Foundation postdoctoral fellowship of the Open Philanthropy Project. G.H and S.L.S. are supported by the Max Planck Society and the Collaborative Research Center 1507 funded by the Deutsche Forschungsgemeinschaft (DFG project number 450648163).

## Author Contributions

The study was designed and conceived by A.G.J. and P.J.K. All cell growth and biochemical assays were performed by A.G.J. Protein purification and detergent screens were performed by A.G.J. and N.K.M.-B. Samples for cryo-EM were prepared by A.G.J. and M.L.M. EM data collection and processing was performed by A.G.J. and M.L.M. Model building and analysis was performed by A.G.J., S.L.S., and P.J.K. Molecular dynamics simulations were performed by S.L.S. and G.H. Figures were prepared by A.G.J., M.L.M., and S.L.S. The manuscript was written by A.G.J. and P.J.K. All authors contributed to editing the manuscript and support the conclusions.

## Competing Interests

None declared.

Correspondence and requests for materials should be addressed to A.G.J. and P.J.K.

## Methods

### Bacterial growth assays

pBAD plasmids encoding full-length and ΔCTD GSDMs were constructed using gene synthesis (IDT) or PCR amplification from existing plasmids followed by Gibson assembly. Constructs expressing single amino acid mutations were generated by “around the horn” (ATH) mutagenesis or gene synthesis (IDT). The following GSDM sequences were used in this study: hGSDMD (Uniprot ID P57764), *Runella* bGSDM (IMG ID 2525253496), Unclassified *Bacteroidetes* metagenomic isolate bGSDM (simplified to “*Bacteroidetes* bGSDM”, IMG ID 2806880301), *Nostoc* bGSDM (IMG ID 2631173059), *Vitiosangium* bGSDM (IMG ID 2831770670), and *Ideonella* bGSDM (IMG ID 2684147428). C-terminal truncations for the ΔCTD GSDMs were based on experimentally demonstrated or predicted cleavage sites. For cleavage site predictions, proteolysis was assumed to occur within exposed loops of X-ray crystal or AlphaFold predicted structures, and before a small P1′ residue (Gly, Ala, or Ser). The ΔCTD GSDMs had the following sequence lengths: 1–275 (hGSDMD), 1–247 (*Runella* bGSDM), 1–247 (*Bacteroidetes* bGSDM), 1–246 (*Nostoc* bGSDM), 1–237 (*Vitiosangium* bGSDM), and 1–244 (*Ideonella* bGSDM). Sequence-verified plasmids were transformed into *E. coli* TOP10 cells, the *E. coli* triple cardiolipin synthase knockout strain BKT12^30^, or the parental *E. coli* strain W3310 used to make BKT12. Transformations were plated on LB agar plates with 100 µg mL^−1^ ampicillin and 1% glucose. Single colonies were used in duplicate or triplicate to inoculate 5 mL liquid cultures in LB with 100 µg mL^−1^ ampicillin and 1% glucose and were incubated with shaking overnight at 37°C. The next morning, cultures were 10-fold serially diluted in 1× phosphate-buffered saline (PBS) from 10^0^ to 10^−7^ and 5 µL of each dilution was spot onto LB agar plates with 100 µg mL^−1^ ampicillin and 1% glucose (to repress expression) or 0.2% arabinose (to induce expression from the arabinose promoter). Spots were allowed to dry at room temperature and plates were then grown overnight at 37°C. The plates were imaged using a Chemidoc system (Bio-Rad) and colony forming units (CFUs) were determined by counting colonies at appropriate dilutions. For conditions in which no colonies appeared in the 10^−1^ and the 10^0^ dilution was uncountable, it was assumed that 1 colony was present in 10^0^ and therefore 200 CFU mL^−1^ total.

### Recombinant protein expression and purification

bGSDM sequences were engineered for controlled activation by inserting site-specific protease cleavage motifs into predicted or verified cleavage sites using the target cleavage sequences for Tobacco Etch Virus (TEV) protease (ENLYFQ/G), the human rhinovirus (HRV) 3C protease (LEVLFQ/GP), the thrombin protease (LVPR/GS), or the WELQut protease (WELQ/G) where the backslash indicates the cleaved bond. Mammalian GSDMs were engineered for controlled activation by HRV 3C as previously described^6,20^, by inserting the cleavage site immediately after E262 in mGSDMA3 or replacing residues 259–275 of hGSDMD with the cleavage site. GSDM-encoding sequences were codon-optimized for *E. coli* and constructed by DNA synthesis (IDT). Gene fragments were cloned by Gibson assembly into a custom pET vector for expression fused to a 6×His-tagged SUMO2 tag. The N-terminal methionine residue was removed from each sequence and replaced by a single serine residue left as a scar following SUMO2 tag removal by the human SENP2 protease. For select constructs, alanine mutants of N-terminal cysteines were prepared by ATH mutagenesis or DNA synthesis (IDT). 40 total engineered bGSDM constructs were screened, and the best identified constructs were for the *Bacteroidetes* bGSDM with the TEV site replacing residues E242–G248, the *Nostoc* bGSDM with the WELQut site replacing residues V243–G247, the *Vitiosangium* bGSDM with the thrombin site replacing residues P234–M239, and the *Ideonella* bGSDM with the WELQut site replacing residues A241–G245.

Plasmids were transformed into BL21(DE3)-RIL *E. coli* and plated on MDG (0.5% glucose, 25 mM Na_2_HPO4, 25 mM KH_2_PO_4_, 50 mM NH_4_Cl, 5 mM Na_2_SO_4_, 2 mM MgSO_4_, 0.25% aspartic acid, 100 mg mL^−1^ampicillin, 34 mg mL^−1^ chloramphenicol, and trace metals) agar plates. 3–5 of the resulting colonies were used to inoculate starter cultures of MDG media for overnight growth at 37°C. MDG starter cultures were used to inoculate 2× 1 L of M9ZB media (0.5% glycerol, 1% Cas-amino Acids, 47.8 mM Na_2_HPO_4_, 22 mM KH_2_PO_4_, 18.7 mM NH_4_Cl, 85.6 mM NaCl, 2 mM MgSO_4_, 100 mg mL^−1^ ampicillin, 34 mg mL^−1^ chloramphenicol, and trace metals) and grown at 37°C with shaking at 230 rpm until the culture reached an OD_600_ of ∼2.5. Cultures were then chilled on ice for 20 min and induced for expression with 0.5 mM IPTG before shaking overnight at 16°C and 230 rpm. The next morning, expression cultures were pelleted by centrifugation, washed with PBS, re-pelleted, and pellets were flash frozen on liquid nitrogen and stored at −80°C until purification.

Protein purification was performed at 4°C with all buffers containing 20 mM HEPES-KOH (pH 7.5) and other components as described below. Cell pellets were thawed and lysed by sonication in buffer containing 400 mM NaCl, 30 mM imidazole, and 1 mM DTT. Lysate was clarified by centrifugation and passing through glass wool, bound to NiNTA agarose beads (QIAGEN), which were subsequently washed with buffer containing 1 M NaCl, 30 mM imidazole, and 1 mM DTT. Protein was eluted with buffer containing 400 mM NaCl, 300 mM imidazole, and 1 mM DTT. Human SENP2 protease was added to the eluate to cleave off the SUMO2 tag before dialyzing overnight into buffer containing 125–250 mM KCl and 1 mM DTT. For initial screening of candidates, the dialyzed protein was directly purified by size-exclusion chromatography (SEC) using a 16/600 Superdex 75 column equilibrated with 250 mM KCl and 1 mM TCEP. For select candidates, SEC was preceded by passing dialyzed protein through a 5 mL HiTrap Q ion exchange column (Cytiva) in 125 mM KCl and 1 mM DTT. Sized proteins were brought to concentrations >20 mg mL^−1^, using 10 kDa molecular weight cut-off concentrators (Millipore), flash frozen on liquid nitrogen, and stored at −80°C until use.

### Liposome assays

Liposomes were prepared from 8 mg lipid total from powder or chloroform dissolved stocks (Avanti) of 1,2-dioleoyl-sn-glycero-3-phosphocholine (DOPC), 1-palmitoyl-2-oleoyl-glycero-3-phosphocholine (POPC), a 3:1 w/w mixture of DOPC or POPC and cardiolipin (CL), or *E. coli* polar lipid extract (*E. coli* liposomes). Lipids were dried by transferring to a glass vial and evaporating chloroform under a stream of nitrogen gas and held under vacuum overnight in the dark. Liposomes for electron microscopy were prepared by resuspending the dried lipids with 20 mM HEPES-KOH (pH 7.5) and 150 mM NaCl. To prepare liposomes for terbium-based liposome rupture assays, and the resuspension buffer was supplemented with 15 mL TbCl_3_ and 50 mM sodium citrate. Liposomes were resuspended in a 800 µL volume (10 mg mL^−1^) by vortexing for 2 min, transferring to a 1.7 mL microfuge tube, and freeze thawing 3× with liquid nitrogen and a warm water bath. The crude liposome suspension was transferred to a glass tube, pulled into a glass syringe (Hamilton), and then extruded 21× with a mini-extruder (Avanti) using 19 mm nucleopore membranes with 0.2 µM pores (Whatman) and 10 mm filter supports (Avanti). Extruded liposomes were purified by SEC with a 10/300 Superdex 200 column equilibrated with 20 mM HEPES-KOH (pH 7.5) and 150 mM NaCl. Fractions containing liposomes were combined and stored at 4°C until use.

Liposome leakage assays were carried out with purified DOPC or *E. coli* liposomes containing TbCl_3_. 40 µL liposome reactions were assembled in a clear well 96-well PCR plate (BioRad) from 24 µL liposomes and 16 µL dipicolonic acid (DPA) dilution buffer at a final concentration of 20 µM DPA. For the data reported in Figure 1, Figure 4, and Extended Data Figure 2, cleavage reactions were assembled in triplicate for the addition of 10 µL per well with final concentrations of 50 µM bGSDM with or without paired proteases in a buffer containing 20 mM HEPES-KOH (pH 7.5) and 150 mM NaCl. Site-specific proteases were used at the following final concentrations: 5 µM homemade TEV protease, 0.1 µM HRV 3C protease, 0.25 units mL^−1^ thrombin protease (MP Bioscience), or 0.1 units µL^−1^ WELQut protease (ThermoFisher). For the data reported in Extended Data Fig. 3 and Extended Data Fig. 11, cleavage reactions were assembled in duplicate in the same buffer with the addition of 10 µL per well with final concentrations of 5 µM bGSDM with or without paired proteases used at the following final concentrations: 1 µM recombinant TEV protease, 0.05 µM human Rhinovirus (HRV) 3C protease, 0.1 units mL^−1^ thrombin protease (MP Bioscience), or 0.05 units µL^−1^ WELQut protease (ThermoFisher). After adding proteases, cleavage reactions were immediately added to wells containing liposomes and plate contents were spun to the bottom of wells by brief centrifugation. The plate was sealed and transferred to a Synergy HI microplate reader (BioTek) to monitor fluorescence resulting from Tb^3+^–DPA complex formation for 6 h with measurements every 2 min by excitation/emission at 276/545 nm.

### Detergent screens and extraction of bGSDM pores

4 mL bGSDM cleavage reactions were prepared in glass tubes with a buffer containing 20 mM HEPES-KOH (pH 7.5) and 150 mM NaCl. Reactions contained final concentration of liposomes at ∼0.8 mg mL^−1^, bGSDMs at 20 µM, and proteases as described above. Screens were initially performed using liposome compositions found to be most ideal by negative-stain EM analysis. Reactions were carried out at room temperature overnight. The next day, liposomes were harvested by centrifuging 150 µL portions at 60,000 × g at 4°C for 30 min with a TLA100 rotor. The supernatant containing uncleaved bGSDM and protease was removed, and pellets were resuspended using 1/5 or 1/10 dilutions of detergents stocks from the JBScreen (Jena Bioscience) or Detergent Screen (Hampton). Detergent resuspensions were incubated for 30 min at room temperature, centrifuged at 60,000 × g at 4°C for 30 min with a TLA100 rotor, and the supernatants were analyzed by SDS-PAGE. Select candidate extractions were analyzed by negative-stain EM and cryo-EM. All extractions were stored at 4°C until use. Of the detergents screened, the *Bacteroidetes* bGSDM pores were found to extract in 0.34 mM DDM, the *Vitiosangium* bGSDM pores were found to extract in 15.6 mM HECAMEG, and the *Vitiosangium* bGSDM slinky-like oligomers were found to extract in 5.16 mM DDMAB. HECAMEG-based extraction of the *Vitiosangium* bGSDM pore was optimized by titrating a range of detergent concentration against the *Vitiosangium* bGSDM incorporated into *E. coli* liposomes.

### Negative-stain electron microscopy

Select bGSDMs were cleaved with paired proteases in the presence of liposomes in a volume of 50 µL in a buffer containing 20 mM HEPES-KOH (pH 7.5) and 150 mM NaCl. Reactions contained liposomes at concentrations of ∼4 mg mL^−1^ and all bGSDMs at 25 µM with proteases concentrations as described above. Samples were incubated at room temperature overnight in glass tubes. To confirm cleavage, samples made in parallel without liposomes were analyzed by 15% SDS-PAGE and stained by Coomassie blue (Extended Data Fig. 2). To identify optimal liposomes for imaging with each bGSDM pore sample, reactions were prepared using liposomes composed of DOPC or POPC lipids with or without CL. Reactions were diluted 1/10 in 20 mM HEPES-KOH (pH 7.5) and 150 mM NaCl buffer and 5 µL was applied to glow-discharged 400 mesh copper grids coated with formvar/carbon film (Electron Microscopy Sciences). Grids were blotted after 1 min, stained with 2% uranyl formate for 2 s, blotted immediately, and air dried. Imaging was performed at 80 kV using the JEOL JEM 1400plus using AMT Image Capture Engine Software version 7.0.0.255. Select samples were imaged at 40,000× magnification to acquire enough micrographs for measuring the inner diameters of >50 pores. For Figure 1 and Extended Data Figure 2, *Bacteroidetes*, *Nostoc*, and *Ideonella* bGSDM pores were all imaged in DOPC liposomes, while the *Vitiosangium* bGSDM pores were imaged from POPC–CL liposomes. For Extended Data Figure 2, *Bacteroidetes* and *Vitiosangium* bGSDM pores were imaged from *E. coli* liposomes. Pore dimensions were measured manually using ImageJ version 2.9.0. Select detergent-extracted bGSDM pore samples were applied to grids and imaged and measured in the same manner as liposomes.

### Cryo-EM sample preparation and data collection

For all bGSDM samples, 3 μL of sample was vitrified on grids using a Mark IV Vitrobot (ThermoFisher). 30 min prior to sample vitrification, grids were glow discharged using an easiGlow^TM^ (Pelco). Grid type and blotting time was optimized for each sample using a double-sided blot with a constant force of 0, in a 100% relative humidity chamber at 4°C, and a 10 s wait prior to plunging in liquid ethane before storing in liquid nitrogen. For the *Bacteroidetes* pores embedded in liposomes, the 1.2/1.3 Carbon Quantifoil^TM^ grids were used with a blotting time of 10 s. For the DDM-extracted *Bacteroidetes* bGSDM pores, lacey carbon grids with carbon thin film support were used with a blotting time of 9 s. For the *Vitiosangium* bGSDM pores and pore-slinky mixtures extracted in HECAMEG detergent, lacey carbon grids with an additional 2 nm layer of carbon thin film were used with a blotting time of 9 s. The pore-slinky mixture represents the same sample as the pore-only sample, after leaving it for one week at 4°C. For the *Vitiosangium* bGSDM oligomeric slinkies extracted in DDMAB detergent, 2/1 Carbon Quantifoil^TM^ grids with an additional 2 nm layer of carbon thin film were used with a blotting time of 9 s. For the HECAMEG-extracted *Vitiosangium* bGSDM pore-slinky mixture that yielded side views, 1.2/1.3 UltrAuFoil^Ⓡ^ grids were used with a blotting time of 7 s.

Cryo-EM data were collected either using a Talos Arctica (ThermoFisher) operating at 200 kV or a Titan Krios (ThermoFisher) microscope operating at 300 kV. Both microscopes were equipped with a K3 direct electron detector (Gatan). SerialEM software version 3.8.6 was used for all collections. Data collection on grids of the *Bacteroidetes* pores embedded in liposomes was performed using a Titan Krios microscope and a total of 4,340 movies were acquired at a pixel size of 0.83 Å, a total dose of 54.5 e− /Å^2^, dose per frame of 1.1e− /Å^2^ at a defocus range of −1 to −2 µm. Grids of the DDM extracted *Bacteroidetes* bGSDM pores were imaged using a Talos Arctica microscope and a total of 5,364 movies were acquired at a pixel size of 1.1 Å, a total dose of 51.7 e− /Å^2^, dose per frame of 1.1e− /Å^2^ at a defocus range of −1.0 to −2.0 µm. Data collection on grids of the *Vitiosangium* bGSDM pores extracted from POPC-CL liposomes in HECAMEG detergent was performed using a Titan Krios and a total of 8,930 movies were acquired at a pixel size of 0.83 Å, a total dose of 48.35 e− /Å^2^, dose per frame of 1.1− /Å^2^ at a defocus range of −1.0 to −2.0 µm. Data collection on grids of the *Vitiosangium* bGSDM slinky-like oligomers extracted from POPC-CL liposomes in DDMAB detergent was performed using a Titan Krios microscope and a total of 8,156 movies were acquired at a pixel size of 1.06 Å, a total dose of 51.8 e− /Å^2^, dose per frame of 1.04 e− /Å^2^ at a defocus range of −0.7 to −2.0 µm. Data collection on grids of the *Vitiosangium* bGSDM pore-slinky mixture extracted from POPC-CL liposomes in HECAMEG was performed using a Titan Krios microscope and a total of 4,818 movies were acquired at a pixel size of 0.83 Å, a total dose of 52.68 e− /Å^2^, dose per frame of 1.1e− /Å^2^ at a defocus range of −1.0 to −2.0 µm. Data collection on grids of the *Vitiosangium* bGSDM pore-slinky mixture extracted from *E. coli* liposomes in HECAMEG was performed using a Titan Krios microscope and a total of 33,411 movies were acquired at a pixel size of 1.3 Å, a total dose of 53 e− /Å^2^, dose per frame of 1.06 e− /Å^2^ at a defocus range of −1.0 to −2.8 µm. Micrographs shown in Figure 2 and Extended Data Fig. 4, 6, and 10 were prepared using IMOD version 4.11.3 and ImageJ Version 2.9.0.

### Cryo-EM image processing and model building

For the *Bacteroidetes* bGSDM pores and the *Vitiosangium* bGSDM pores and pore-slinky mixtures, movies frames were imported to cryoSPARC^34^ for patch-based motion correction, patch-based CTF estimation, and 2D classification. For the *Vitiosangium* bGSDM oligomeric slinkies, movies frames were pre-processed using the on-the-fly-processing scheme of RELION (version 3.1)^35^. Motion correction was performed using MotionCor2^36^ and the motion corrected movies were imported to cryoSPARC for patch-based CTF estimation, 2D and 3D particle classification, non-uniform refinement, and masked local refinement. The resulting cryo-EM density was post-processed using DeepEMhancer^37^ prior to model building. For the *Vitiosangium* bGSDM pore-slinky mixture yielding pore sideviews, movies frames were imported to cryoSPARC^34^ for patch-based motion correction, patch-based CTF estimation. Particles were picked using the Topaz^38^ wrapper inside cryoSPARC before extraction and 2D classification, 3D classification, and homogenous and masked local refinements (see Extended Data Fig. 10 for processing workflow). Local resolution was estimated using cryoSPARC and visualized in ChimeraX.

The globular domain of the inactive-state *Vitiosangium* bGSDM (7N51) was used as a starting model after docking it into a single protomer of the slinky-like oligomer using Coot^39^. A complete protomer model was manual built in Coot and iterative corrections were made based on Phenix real-space refine^40^. The protomer model was validated in Phenix with MolProbity^41^. The 54-mer slinky model in Fig. 3d was constructed by extending from a docked protomer by continuously fitting new protomers within the cryo-EM map density. To make the 52-mer pore model in Fig. 3d, the protomer models were realigned using a custom script and a geometric model based on the radius of a low-quality 3D reconstruction of anisotropic cryo-EM data of the 52-mer pore. Protomers were realigned to preserve the inter-subunit hydrogen bonding pattern of the slinky-like oligomer. Structure figure panels were generated using UCSF ChimeraX^42^ and PyMOL version 2.4.0.

### Molecular dynamics simulations

*Membrane.* We set up the systems for molecular dynamics (MD) simulations using Charmm-GUI^43^. As membrane template, we used 15.2 × 15.2 nm^2^ patch mimicking the bacterial inner membrane (PVPE:PVPG:cardiolipin, 15:4:1) surrounded with 150 mM NaCl solution. We minimized the potential energy of this patch in 5,000 steepest descent steps and performed 250 ps of NVT and 1.625 ns of NPT equilibration while stepwise decreasing the initial positional restraints of the phosphate z positions and the lipid tail dihedral angles. We used this membrane to set up simulation systems of 1–3 subunits of the *Vitiosangium* bGSDM. For simulations of the full 52-mer pore we multiplied the membrane patch along the xy plane to create a 62.3 × 62.3 nm^2^ large membrane patch.

*bGSDM.* We set up 1, 2, 3 and 52-mer *Vitiosangium* bGSDM systems in pore conformation by extracting the respective oligomer structures from the full 52-mer pore, with and without palmitoylated C4, inserting them into the membranes and removing all lipids with atoms overlapping with any protein (or C4 palmitoyl) atom, using Charmm-GUI to create simulation topologies. We alleviated membrane asymmetries by removing lipids from the overfilled leaflet. Subsequently, we resolvated the system with TIP3P water^44^, removed water molecules in the membrane, and then added NaCl to a sodium concentration of 150 mM and chloride atoms to neutralize the charge of the system.

For simulations of the inactive conformation of *Vitiosangium* bGSDM, we removed the 29 aa CTD from the inactive crystal structure (PDB ID 7N51^15^) and directly placed the inactive ΔCTD on the membrane patch. We used this system as starting point for 3 replicate 2-µs room-temperature MD simulations. One of these was then continued in 24 replicate 1-µs MD simulations of 1 µs each at 97°C, each with different inintial velocities. We also performed 24 1-µs simulations of ΔCTD dimers at 97°C. As starting dimer model, we placed two subunits from the last frame of a room-temperature replicate next to each other as in the EM ring structure and resolvated the system. To alleviate pressure in the protein-bound leaflet at elevated temperature, we randomly removed 10 (20) PVPE lipids from the protein-bound leaflet in the monomer (dimer) simulations. The dimer 1-µs production runs were preceded by potential energy minimization, as described below, and equilibration for 50 ns with position restraints on the protein, as described in Supplementary Table 1.

All MD simulations were performed using Gromacs version 2022.4^45^ and the CHARMM36m force-field^46^. We used a standard integration timestep of 2 fs. Electrostatic interactions were computed using the particle-mesh Ewald (PME) algorithm^47^. Real-space electrostatic and van-der-Waals interactions were cut-off at 1.2 nm. We constrained the distance of bonds with hydrogen atoms using LINCS^48^. We maintained a constant temperature of 37°C using the velocity-rescale algorithm^49^ with a time constant of 1 ps, controlling protein, membrane and solvent (water and ions) individually. During system equilibration, we used the Berendsen barostat^50^ for semi-isotropic pressure coupling (x and y dimensions coupled together) with a time constant of 5 ps, a reference pressure of 1 bar, and a compressibility of 4.5 × 10^−5^ bar^−1^. After the equilibration phase, we switched to the Parrinello-Rahman algorithm^51^ for pressure control, while keeping all other control parameters the same.

Each system was first energy minimized for 5,000 steepest descent steps. Then, equilibration simulations of 50 ns length each were performed. During these equilibration simulations, the initial positional restraints of the heavy atoms of the protein (and the phospholipids) were gradually lifted (Supplementary Table 1). No positional restraints were used during the production simulations. Simulations were run for times indicated in Supplementary Table 2 and in the case of replicate simulations were started with randomly drawn initial velocities according to the Maxwell-Boltzmann distribution. The three simulations of the membrane bound inactive structure were not started from the exact same system, but from two systems with initially slightly different conformations of the palmitoyl fatty acid tail.

To calculate the Cα root mean squared deviation (RMSD) of the pore, we first calculated the average root mean squared fluctuation (RMSF) of each residue’s Cα and excluded the highly flexible protein regions (termini and fingertip regions with RMSF > 3.5 Å) from the subsequent RMSD calculation. The RMSD was calculated with respect to the average structure of the last 900 ns of the production simulation.

Two residues on neighboring subunits were considered to form a contact if at least one pair of heavy atoms was within 3.6 Å distance. We estimated the probability of interfacial contacts over the last 900 ns of the simulation of the palmitoylated 52-mer pore and the 52 distinct interfaces.

Visual analysis as well as image and movie rendering were performed using PyMOL, VMD^52^, and UCSF ChimeraX^42^. Quantitative analyses and system setup were implemented with python 3.9 and use the MDAnalyis package (v2.4.2)^53,54^.

## Data Availability Statement

Coordinates and density maps of the active-state *Vitiosangium* bGSDM are being deposited with the PDB and the EMDB under the accession codes 8SL0 and EMD-40570. Simulation parameter files, raw trajectories and code used for analysis are deposited in two zenodo repositories (10.5281/zenodo.7828403, 10.5281/zenodo.8272143). All other data are available in the manuscript or the supplementary materials.

## Code Availability Statement

Custom scripts used to make the *Vitiosangium* bGSDM 52-mer pore model are available in a zenodo repository (10.5281/zenodo.7828403).

---

**Extended Data Fig. 1.**
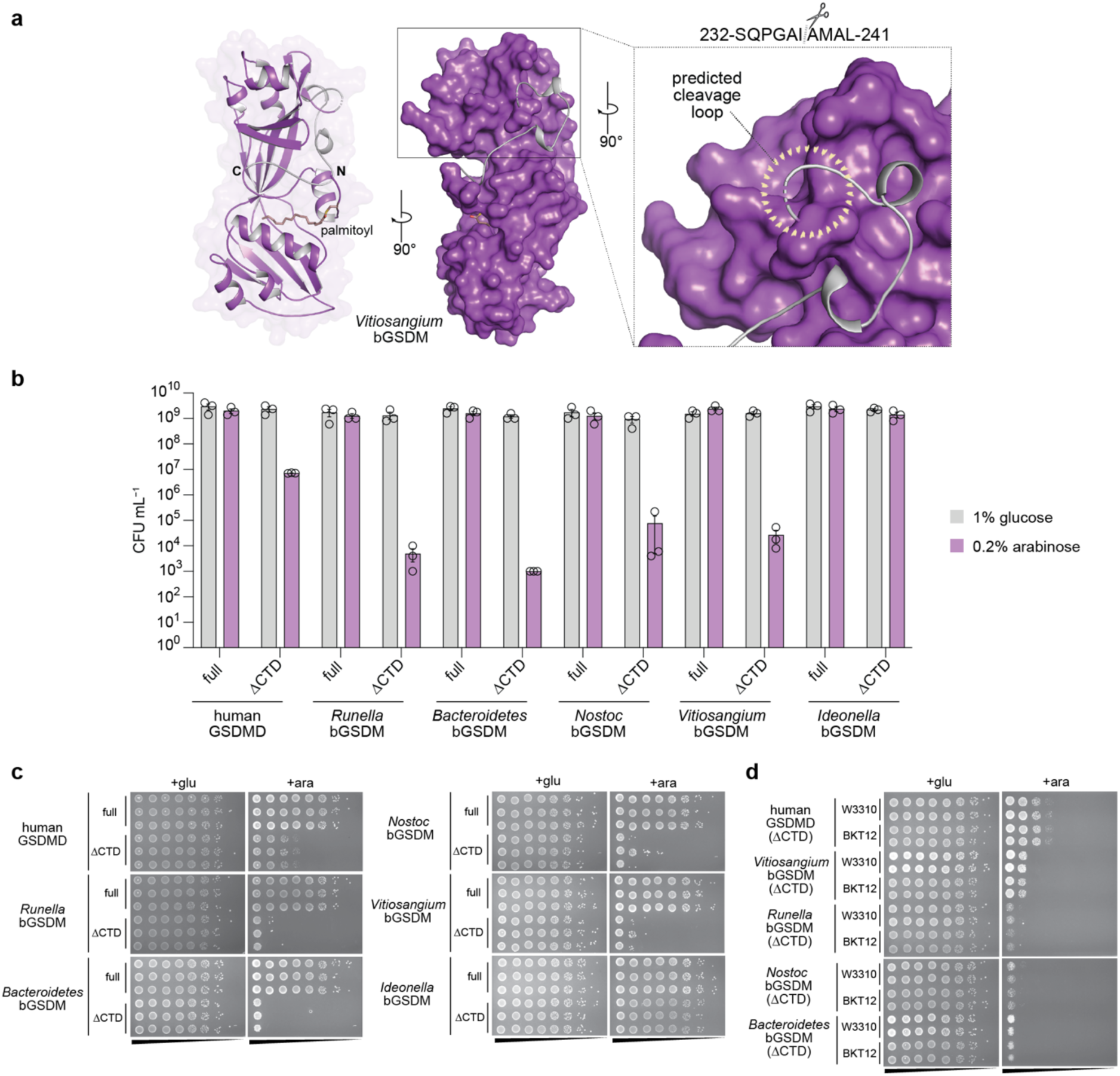
Expression of bGSDM NTDs induces potent cellular toxicity. **a,** Crystal structure of an inactive bGSDM from a *Vitiosangium* species (PDB ID 7N51) and indication of the disordered loop that was targeted for cleavage site engineering. **b,** Colony forming units (CFU) per mL of *E. coli* derived from spot assays shown in panel (c). **c,** TOP10 *E. coli* harboring plasmids encoding full-length GSDMs (full) or the N-terminal pore-forming domain alone (ΔCTD) were grown on LB-agar plates in triplicate. LB-agar contained either 1% glucose or 0.2% arabinose to repress or induce expression, respectively. Cells were serially diluted and plated out from left (10^0^) to right (10^−7^) with 5 µL per spot. Though the *Ideonella* ΔCTD construct does not drastically reduce the CFUs compared to the full construct, colonies grow more slowly and appear fainter in agreement with toxicity from pore formation. **d,** Parental (W3310) or the triple cardiolipin synthase knockout (BKT12) E. coli harboring plasmids for the ΔCTD GSDMs were grown and spotted out onto LB agar plates with 1% glucose or 0.2% arabinose in duplicate as in panel (c).

**Extended Data Fig. 2.**
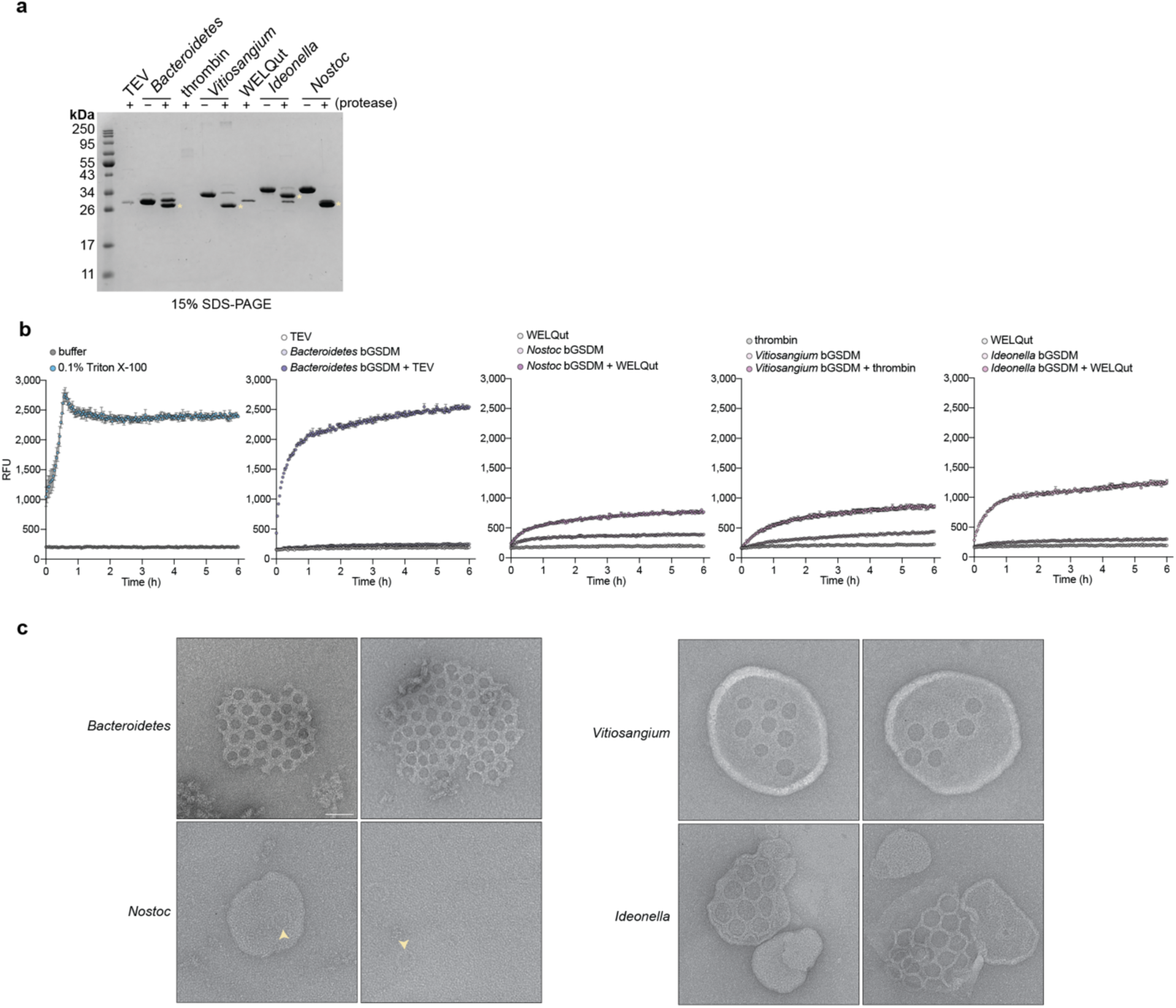
Cleavage site engineering enables analysis of diverse bGSDM pores. **a,** Engineered bGSDMs were treated with or without paired site-specific proteases for 18 h at room temperature and analyzed by 15% SDS-PAGE and visualized by Coomassie staining. Cleaved bGSDM proteins are indicated with a yellow asterisk. **b,** Full time-course of liposome leakage assays related to Fig. 1b. Error bars represent the SEM of three technical replicates. c, Negative-stain EM micrographs representing larger view fields micrographs shown in Fig. 1d or second example micrographs used to measure pore sizes for Fig. 1c. Scale bar = 50 nm.

**Extended Data Fig. 3.**
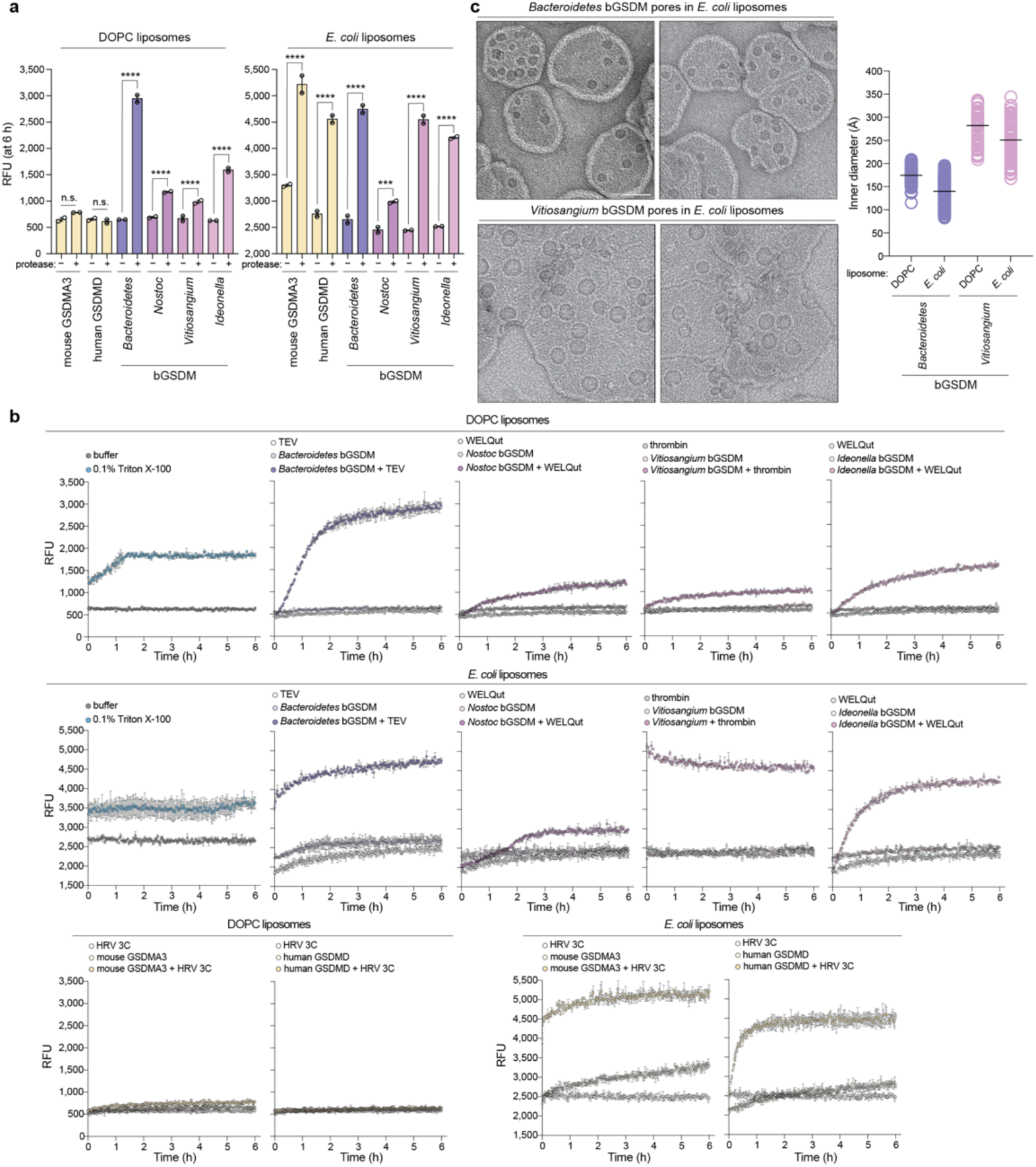
Engineered bGSDMs form pores in liposomes with simple and complex lipid compositions. **a,** Liposome leakage assay of engineered mammalian GSDMs and bacterial GSDMs with matched site-specific proteases. The left plot shows the results from an experiment performed with liposomes prepared from DOPC lipids (DOPC liposomes), while the right plot shows the result from an experiment using liposomes prepared from *E. coli* polar lipid extract (*E. coli* liposomes). The species and/or paralog of mammalian GSDMs or bGSDMs or mammalian GSDMs (and protease sites) are as follows: mGSDMA3 (HRV 3C site), hGSDMD (HRV 3C site), *Unclassified Bacteroidetes* (TEV site), *Nostoc sp. Moss4* (WELQ site), *Vitiosangium sp. GDMCC 1.1324* (thrombin), and *Ideonella sp. 201-F6* (WELQ). Error bars represent the SEM of two technical replicates. n.s. ≥ 0.05; \*\*\**P* < 0.001; \*\*\*\**P* < 0.0001. **b,** Full time-course of liposome leakage assays related to panel (a). Error bars represent the SEM of two technical replicates. **c,** Negative-stain EM micrographs of *Bacteroidetes* bGSDM and *Vitiosangium* bGSDM pores in *E. coli* liposomes (left) and plot comparing pore inner diameters of these bGSDMs in DOPC liposomes versus the same bGSDMs in E. coli liposomes (right). The number of pores measured (n) for each species in *E. coli* liposomes is *Bacteroidetes* (n = 171) and *Vitiosangium* (n = 100). The inner diameters values for pores in DOPC liposomes are the same as in Figure 1c. The black bar represents the average inner diameter of measured pores. Scale bar = 50 nm.

**Extended Data Fig. 4.**
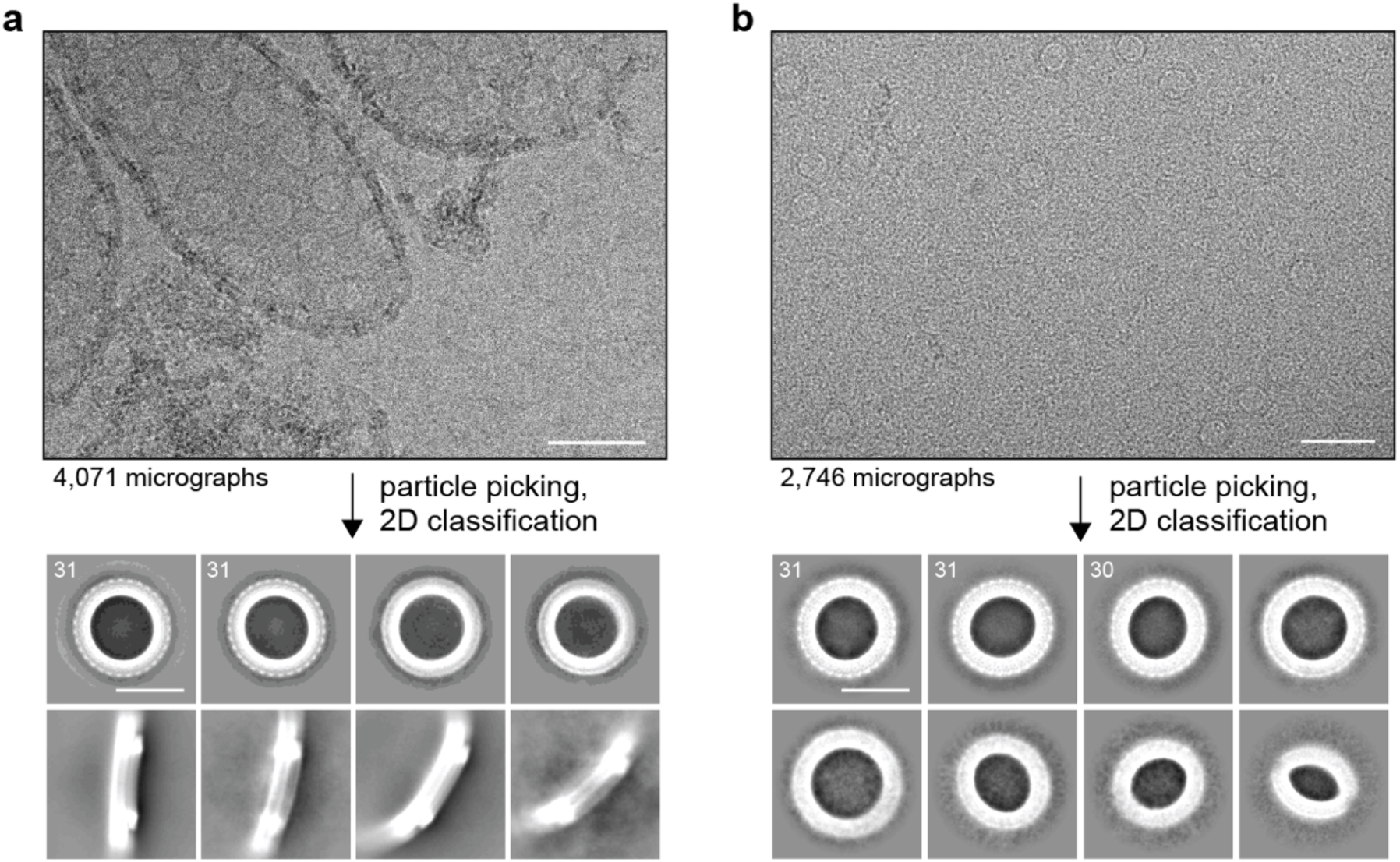
Cryo-EM data processing of *Bacteroidetes* bGSDM pores. **a,** Representative cryo-EM micrograph and select 2D class averages of bGSDM pores from pore-liposome samples. **b,** Representative cryo-EM micrograph and select 2D class averages of DDM-extracted bGSDM pores. Numbers in the upper left-hand corner of 2D classes represent the number of bGSDM protomers observed in that class. Scale bars = 20 nm.

**Extended Data Fig. 5.**
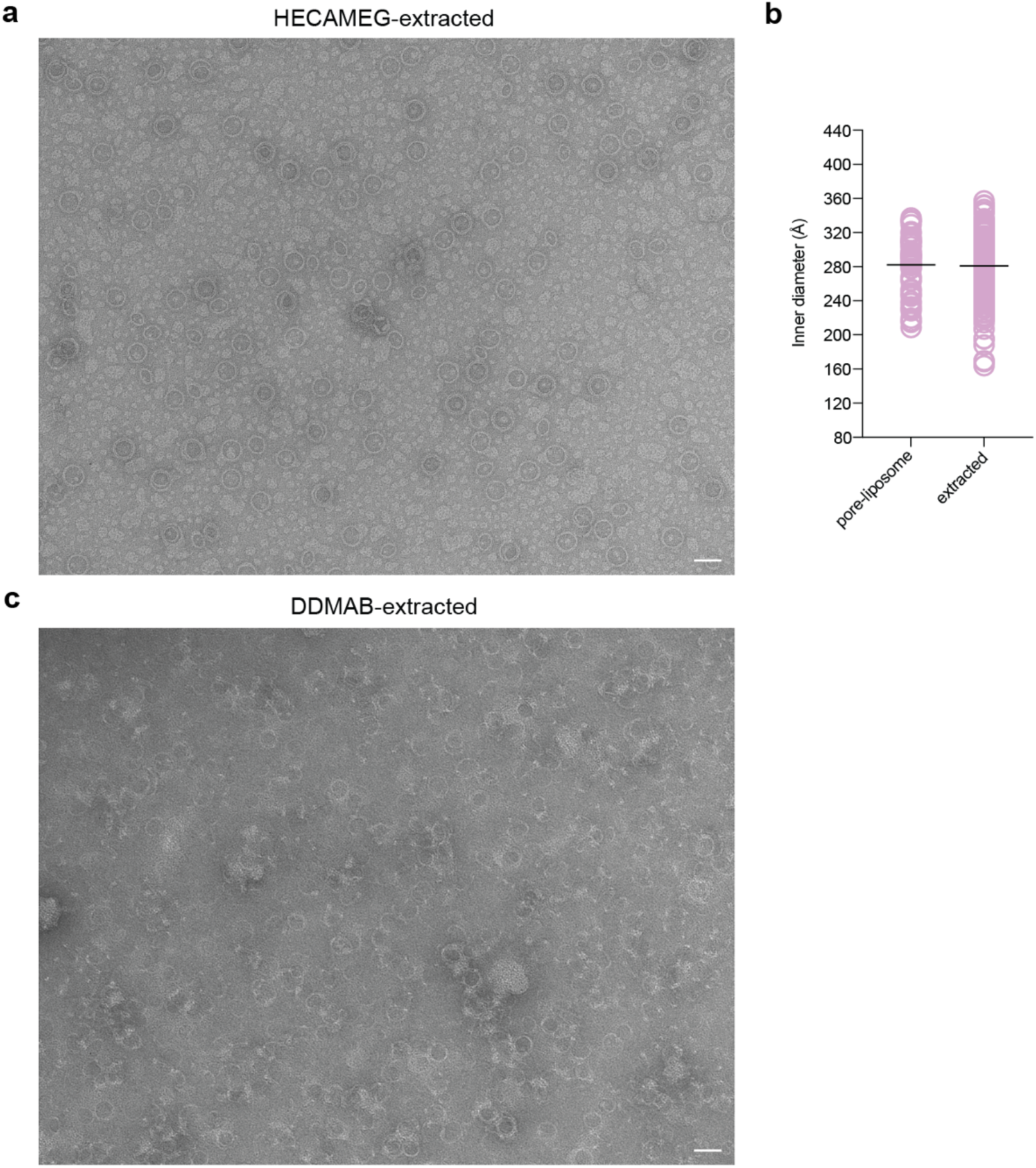
Negative-stain EM micrographs of detergent-extracted *Vitiosangium* bGSDM pores and slinkies. **a,** HECAMEG detergent-extracted bGSDM pores. Scale bar = 50 nm. **b,** Comparison of inner diameters measured from pore-liposome samples (Fig. 1 and Extended Data Fig. 2) and HECAMEG detergent-extracted bGSDM pores. The number of pores measured (n) from each sample is as follows: pore-liposome (n = 56), extracted pore (n = 189). The black bar represents the average inner diameter of measured pores. **c,** DDMAB detergent-extracted bGSDM slinkies. Scale bars = 50 nm.

**Extended Data Fig. 6.**
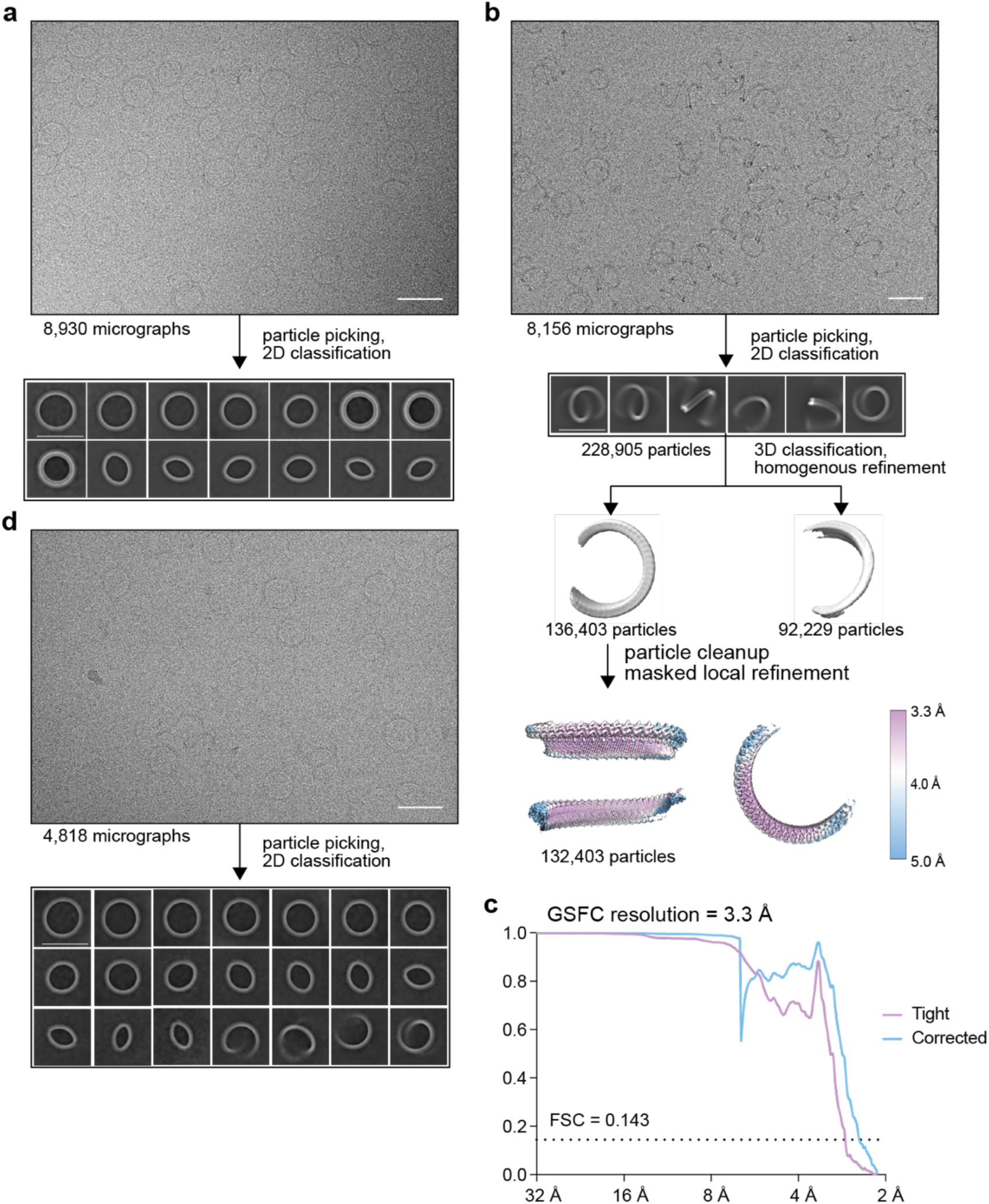
Cryo-EM data processing of *Vitiosangium* bGSDM pores and slinkies. **a,** Representative cryo-EM micrograph and 2D class averages of HECAMEG detergent-extracted bGSDM pores. **b,** Single-particle processing schematic of DDMAB detergent-extracted bGSDM slinkies. From top to bottom: representative cryo-EM micrograph and 2D class averages (as in Fig. 2b), particle classification and map refinement, and local resolution estimate of final map. **c,** Fourier shell correlation (FSC) curves versus resolution of bGSDM slinky map. Resolution was estimated at an FSC of 0.143. Scale bars = 50 nm. **d,** Representative cryo-EM micrograph and 2D class averages of HECAMEG detergent-extracted bGSDM pore-slinky mixture (as in Fig. 2a).

**Extended Data Fig. 7.**
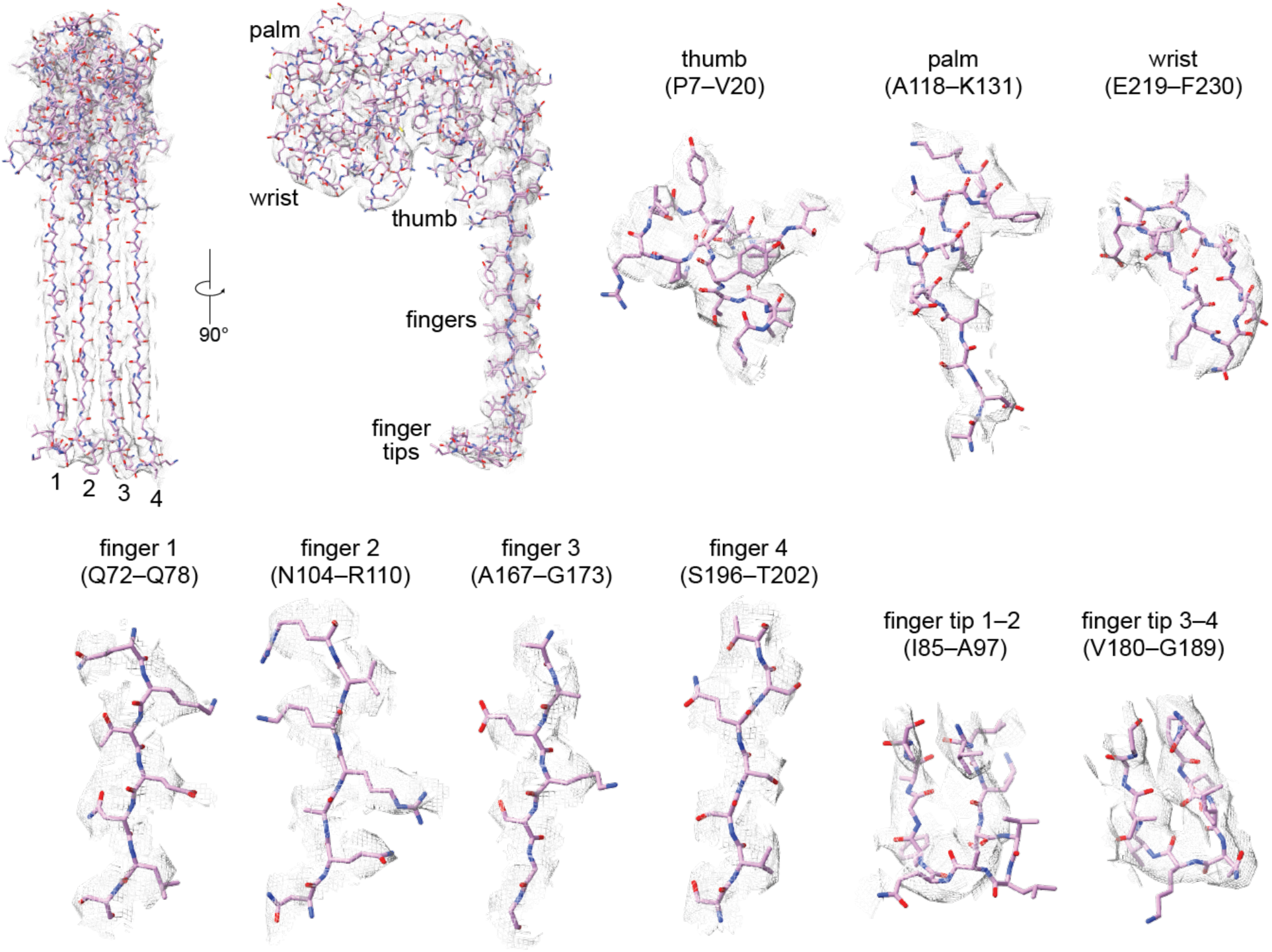
Cryo-EM model to map fitting of bGSDM slinky and pore. **a,** 54-mer model of the *Vitiosangium* bGSDM in a slinky-like oligomerization. **b,** Examples regions of model to map fit quality for a single *Vitiosangium* bGSDM protomer. The map surface has been contoured to 16σ.

**Extended Data Fig. 8.**
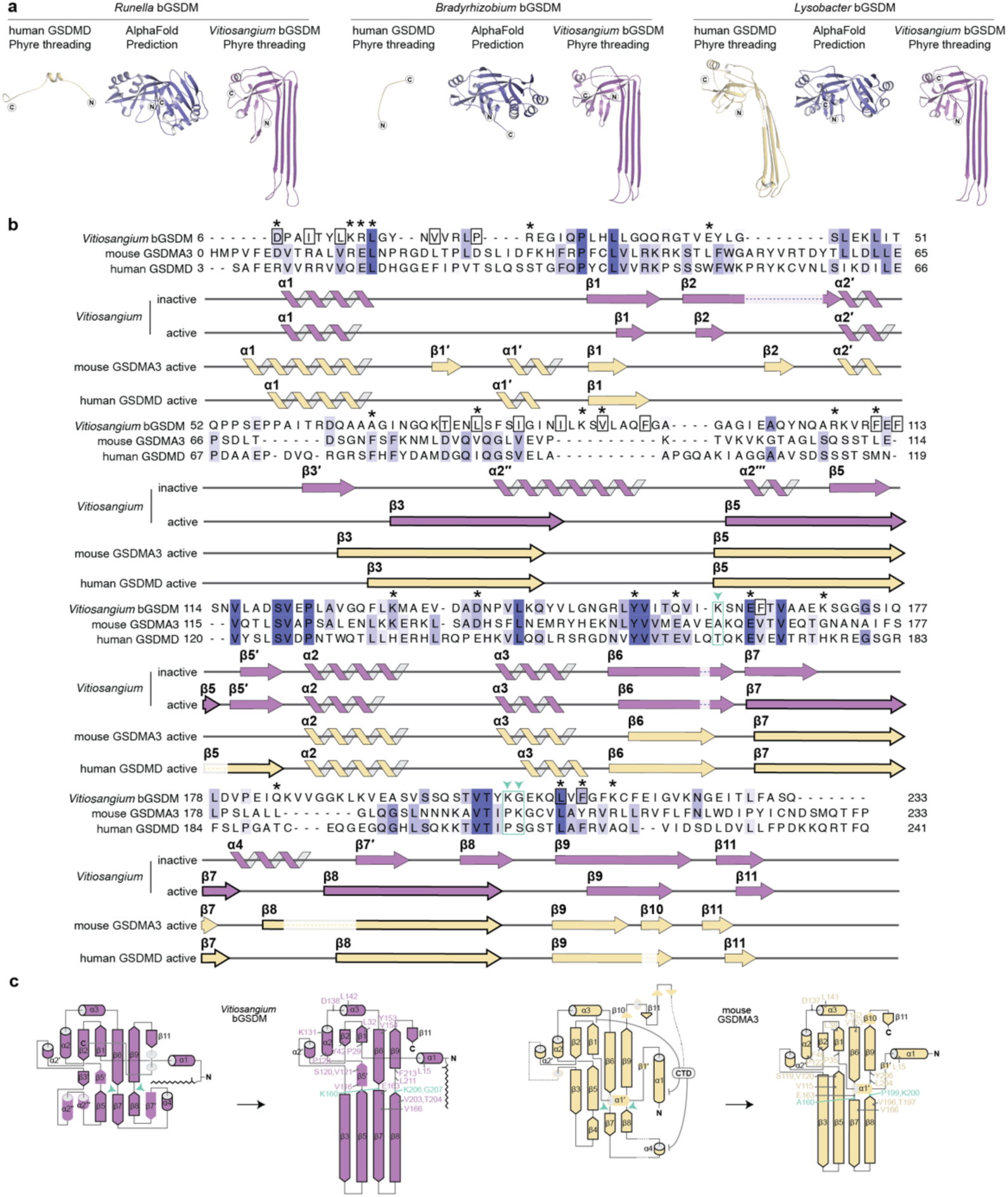
Functional conservation of the GSDM active state. **a,** Predicted bGSDM structures are organized from left to right based on percent sequence identity to the *Vitiosangium* bGSDM: *Runella* bGSDM (18%), *Bradyrhizobium* bGSDM (19%), and *Lysobacter* bGSDM (28%). Phyre homology model threading utilized the active hGSDMD structure (PDB ID 6VFE, left model) or the active *Vitiosangium* bGSDM structure (this study, right model). Each bGSDM structure was also predicted using AlphaFold after deleting ∼20 amino acids from the C-termini and yielded inactive-like structures (center). The modeled sequences are as follows: *Runella* bGSDM (1–247), *Bradyrhizobium* bGSDM (1–237), and *Lysobacter* bGSDM (1–240). **b,** Structure-based alignment of bacterial and mammalian gasdermins. The *Vitiosangium* bGSDM active structure was aligned to the active mGSDMA3 (PDB ID 6CB8) and hGSDMD (PDB ID 6VFE). Secondary structures are indicated below the sequences for each structure, in addition to secondary structures from the *Vitiosangium* bGSDM inactive state crystal structure. Residues of the *Vitiosangium* bGSDM that surround the palmitoyl in the inactive state are boxed in black, asterisks indicate residues that have been mutated to test their effect on bGSDM-mediated bacterial cell death, and green boxes indicate resides that align with the conserved glycines in MACPF/CDC proteins. **c,** Topology diagrams showing the transitions from inactive to active structures of the *Vitiosangium* bGSDM (left) and mGSDMA3 (right). Conserved α-helices and β-strands are outlined in black, the positions of residues universally conserved in charge, identity, or aromaticity are indicated in active state topology diagrams, and green arrows indicate the sites of conserved glycines in MACPF/CDC protein structures.

**Extended Data Fig. 9.**
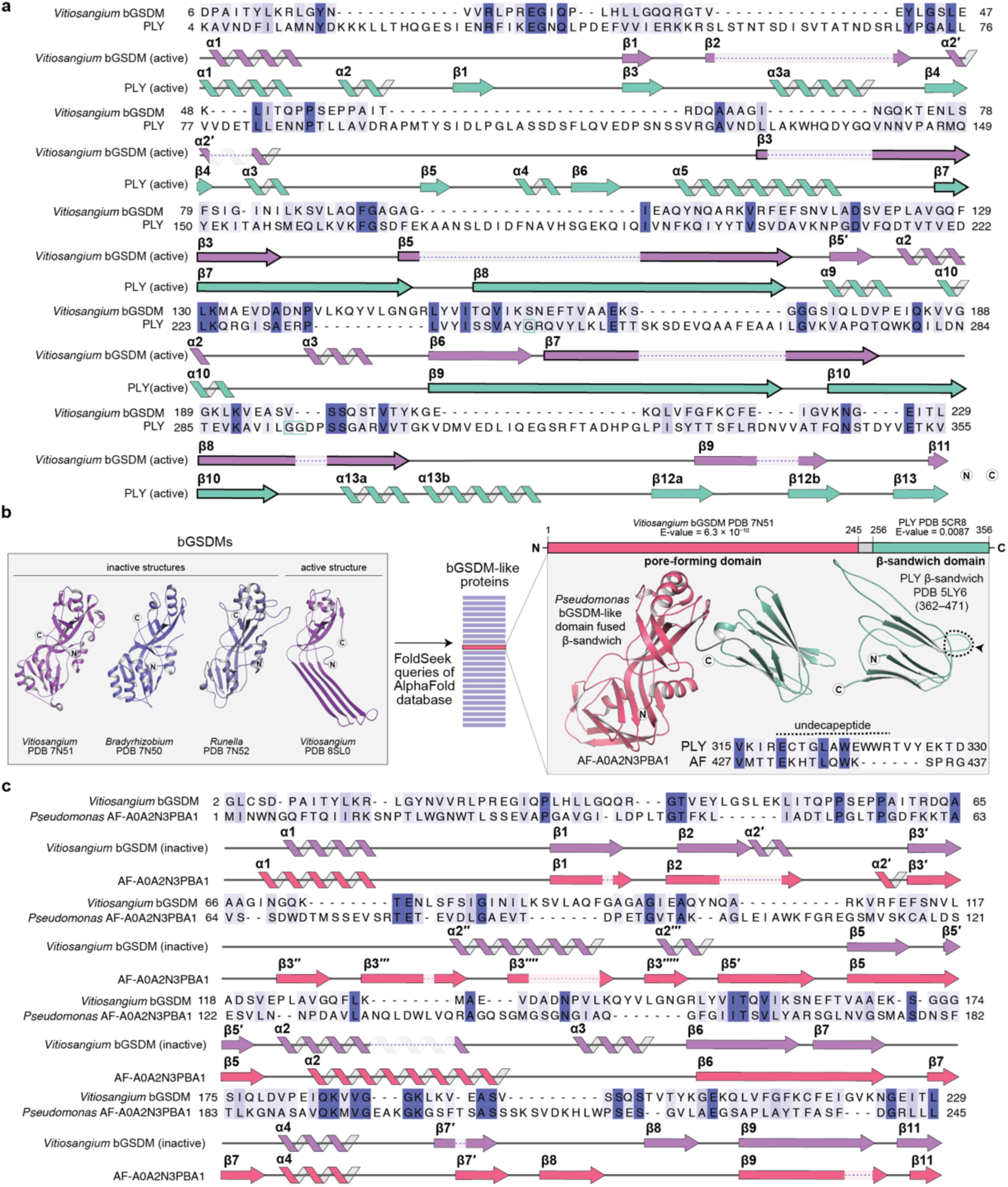
Evidence for a possible divergent evolution of gasdermins and cytolysins. **a,** Structure-based alignment of the *Vitiosangium* bGSDM (aa 6–229) and the pneumolysin (PLY) pore-forming domain (aa 4–355). The *Vitiosangium* bGSDM active state structure (this study, PDB 8SL0) was aligned to the active state PLY structure (PDB ID 5LY6). Secondary structures are indicated below the sequences for each structure and numbered according to prior conventions^19^. Conserved glycine residues of the PLY structure that are present in other MACFP/CDC proteins are boxed in green. **b,** A query of the AlphaFold database with experimentally determined bGSMD structures yielded putative bGSDM-like proteins with cytolysin-like features. A representative structure from a *Pseudomonas* species is shown on the right, indicating the N-terminal bGSDM-like domain in salmon color and the C-terminal immunoglobulin-like β-sandwich domain in green color with similarity to the membrane binding domain of PLY and other cytolysins. The sequence alignment shows the highly conserved undecapeptide present in the β-sandwich domains of multiple cytolysins **c,** Structure-based alignment of the *Vitiosangium* bGSDM (aa 2–229) and the *Pseudomonas* bGSDM-like protein pore-forming domain (aa 1–245). The *Vitiosangium* bGSDM inactive state structure (PDB 7N51) was aligned to the putative inactive state structure of the bGSDM-like protein (AF-A0A2N3PBA1). Secondary structures are indicated below the sequences for each structure, using α-helix and β-sheet numbering for the bGSDM-like protein that reflect homology to the bGSDM.

**Extended Data Fig. 10.**
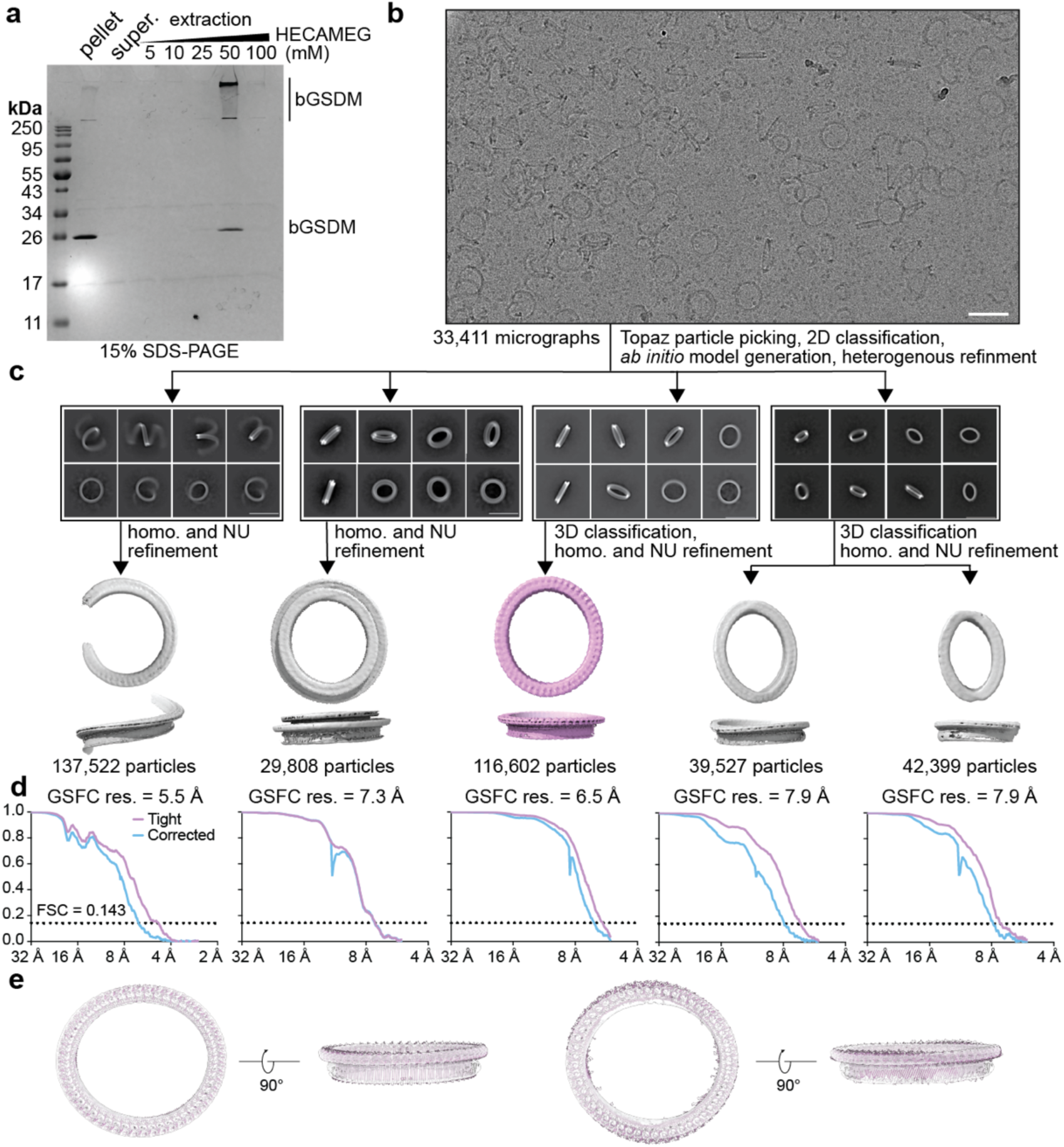
Extraction and cryo-EM data processing of *Vitiosangium* bGSDM pores with sideviews. **a,** Fractions from the detergent extraction of *Vitiosangium* bGSDM pores from *E. coli* liposomes analyzed by SDS-PAGE and Coomassie staining. The bGSDM sample extracted at 50 nm HECAMEG was subsequently used for cryo-EM analysis **b,** Representative cryo-EM micrograph of HECAMEG detergent-extracted bGSDM pores from (a). Scale bar = 50 nm. **c,** 2D class averages from processing micrograph indicated in (b) and single-particle processing schematic of HECAMEG detergent-extracted bGSDM closed-ring pores and slinky-like oligomers. Scale bars = 50 nm. Data was processed using homogeneous (homo.) or non-uniform (NU) refinements. **d,** Fourier shell correlation (FSC) curves versus resolution of bGSDM slinky map. Resolution was estimated at an FSC of 0.143. **d,** Docking of the 52-mer elliptical pore-model into the 6.5 Å resolution map from (c). The left model represents a geometric model based on the structure of the slinky-like oligomer, with an eccentricity of 0.86. The right model represents the maximum eccentricity pore undulation observed in during the MD simulations of the 52-mer pore, with an eccentricity of 0.67.

**Extended Data Fig. 11.**
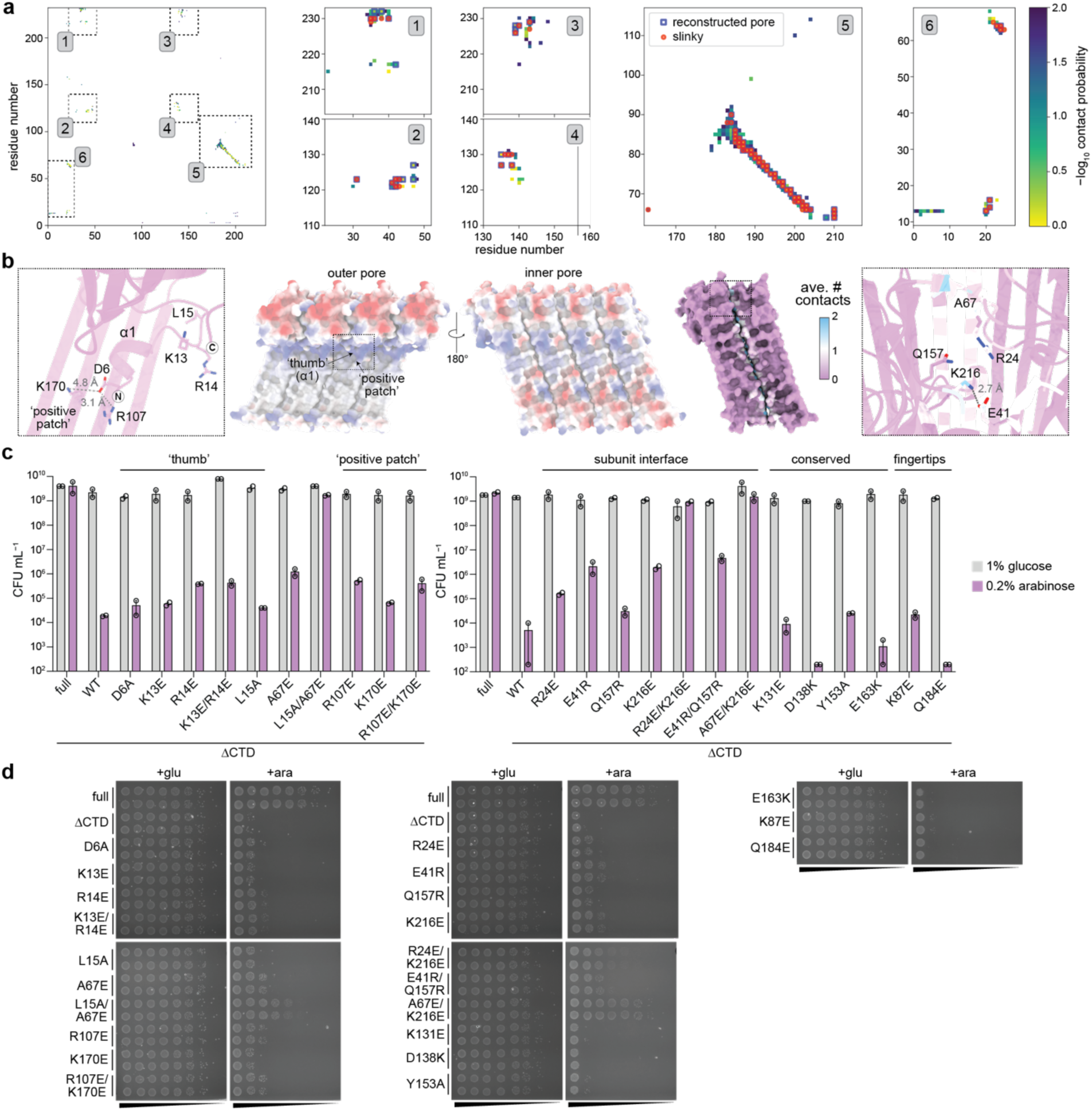
Cell death and liposome rupture by the *Vitiosangium* bGSDM is robust to single mutations. **a,** Residue-residue contacts between neighboring subunits occurring with a frequency of >0.01 over the last 900 ns of an MD simulation of the 52-mer pore with C4-palmitoylation. **b,** Structural representations of the *Vitiosangium* bGSDM oligomer. Center, an electrostatic charge model of a 4-mer of the slinky-like oligomer. Left inset, sites targeted for mutation on the ‘positive patch’ and ⍺1 thumb on the pore exterior. Right, dimer of the Vitiosangium bGSDM along the interface shaded to indicate the frequency of contact occurring over the course of the MD simulation described in panel (a). Right inset, residues targeted for mutagenesis at the subunit interface. **c,** Colony forming units (CFU) per mL of *E. coli* derived from spot assays shown in panel (d). Growth assays test single charge-swap mutations to residues at select interfaces in the active model. **d,** *E. coli* harboring plasmids encoding full-length GSDMs (full) or the N-terminal pore-forming domain alone (ΔCTD) were grown on LB-agar plates in duplicate. LB-agar contained either 1% glucose or 0.2% arabinose to repress or induce expression, respectively. Cells were serially diluted and plated out from left (10^0^) to right (10^−7^) with 5 µL per spot.

**Extended Data Fig. 12.**
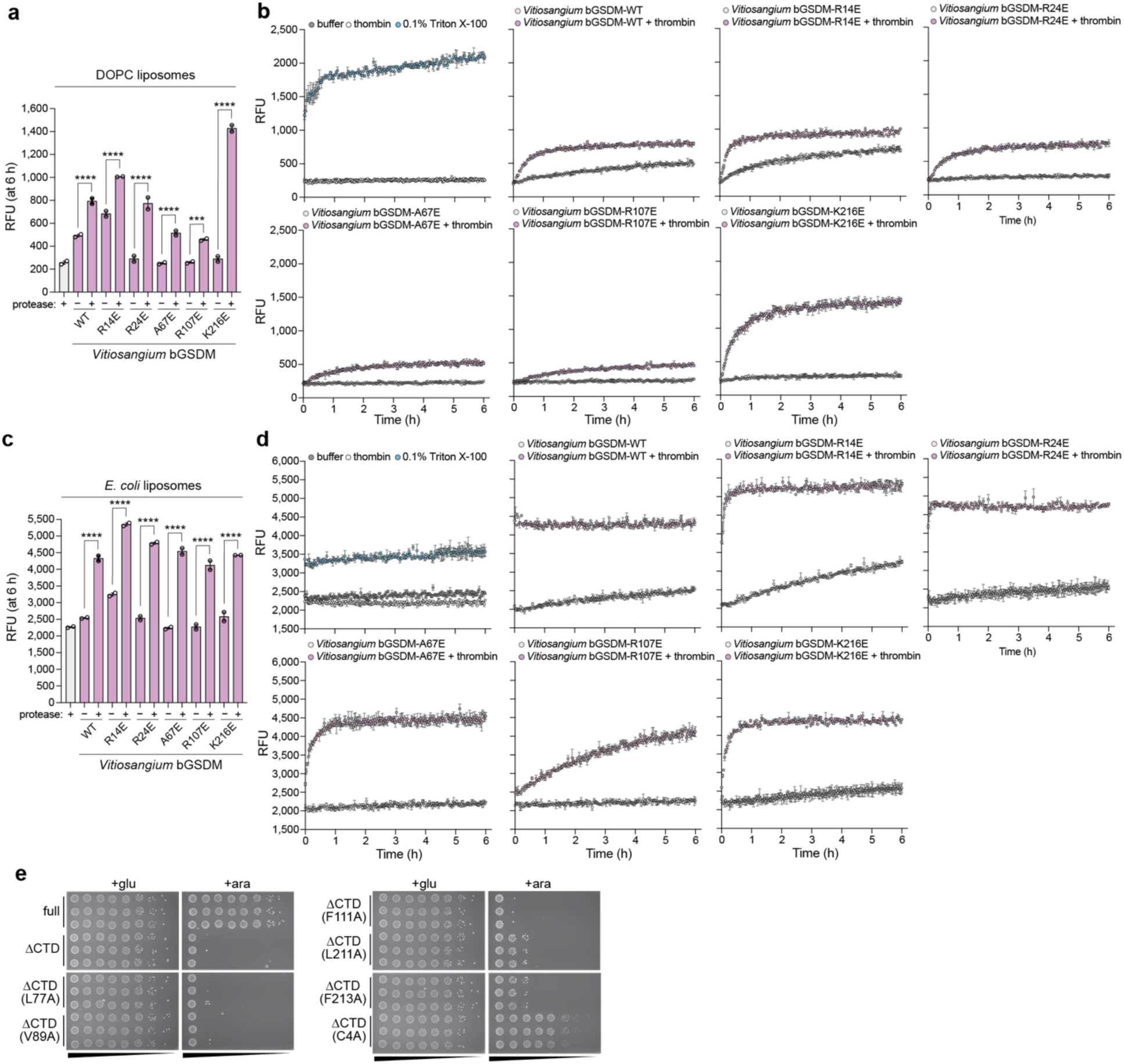
Liposome rupture and cell death by the *Vitiosangium* bGSDM is sensitive to mutation of the palmitoylated cysteine but not other single amino acid residues. **a,** Relative fluorescent units (RFUs) at six hours from liposome rupture experiment testing single amino acid mutants of the *Vitiosangium* bGSDM with DOPC liposomes. **b,** Full time-course RFU data for the plot shown in panel (a). **c,** RFU at six hours for liposome rupture experiment testing single amino acid mutants of the *Vitiosangium* bGSDM with DOPC liposomes. **d,** Full time-course RFU data for the plot shown in panel (c). **e,** Bacterial spot assays testing mutation of residues proximal to the N-terminal cysteine of the Vitiosangium bGSDM. TOP10 *E. coli* harboring plasmids encoding full-length GSDMs (full) or the N-terminal pore-forming domain alone (ΔCTD) were grown on LB-agar plates in triplicate. LB-agar contained either 1% glucose or 0.2% arabinose to repress or induce expression, respectively. Cells were serially diluted and plated out from left (10^0^) to right (10^−7^) with 5 µL per spot.

**Extended Data Fig. 13.**
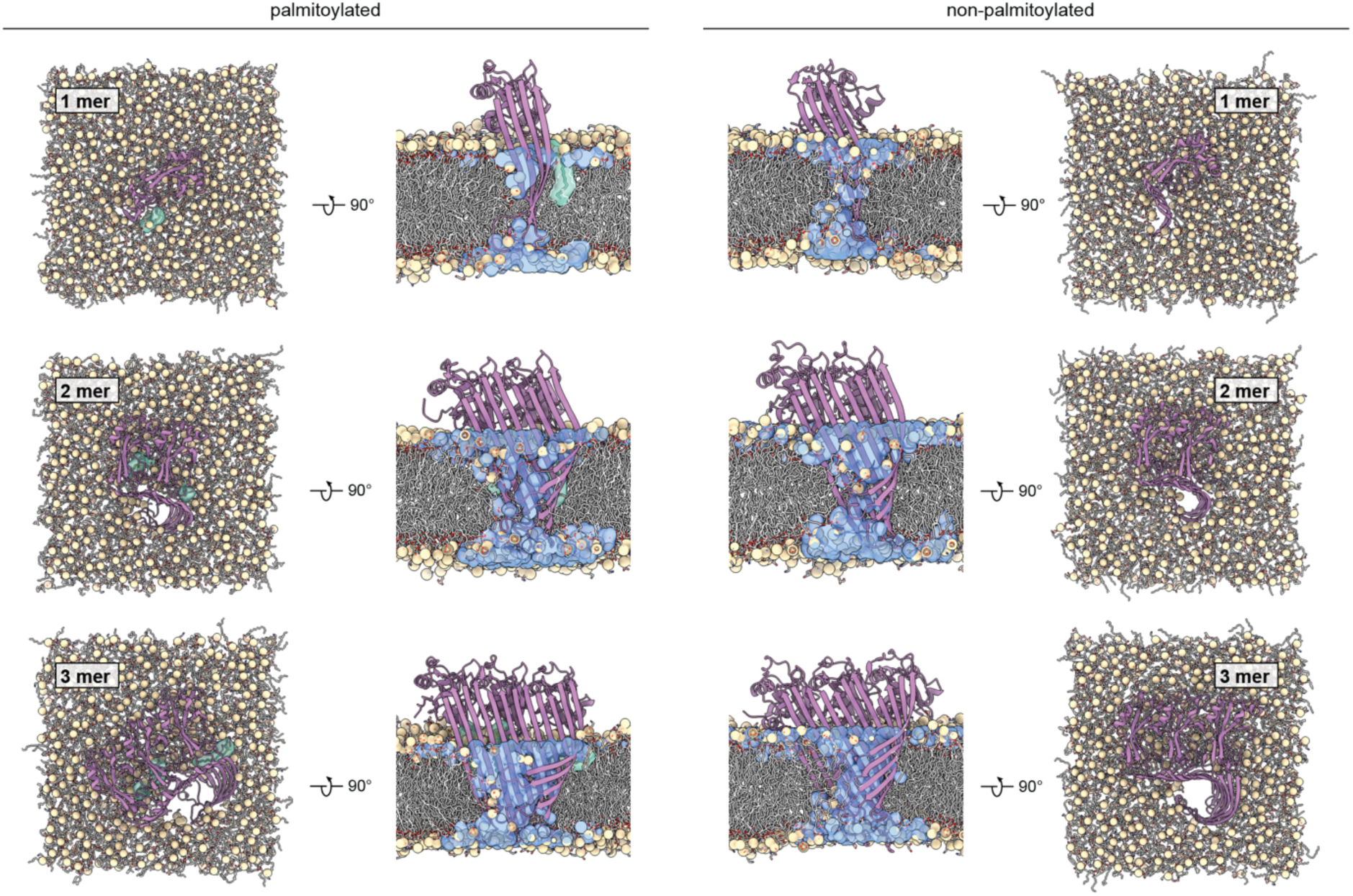
MD simulations of small membrane pores formed by active 1, 2, and 3-mer bGSDM assemblies. Snapshots of palmitoylated (left) and non-palmitoylated (right) mono- and oligomers after 3.3 μs of simulation in a bacterial membrane in top view and side view. Protein shown in purple cartoon representation, membrane phosphates shown as tan spheres, palmitoylated C4 shown in cyan opaque licorice and transparent surface representation. In the side views, water inside the small pores is shown in blue transparent surface representation. Otherwise, solvent molecules and lipid fatty acid tails are omitted for clarity.

**Extended Data Fig. 14.**
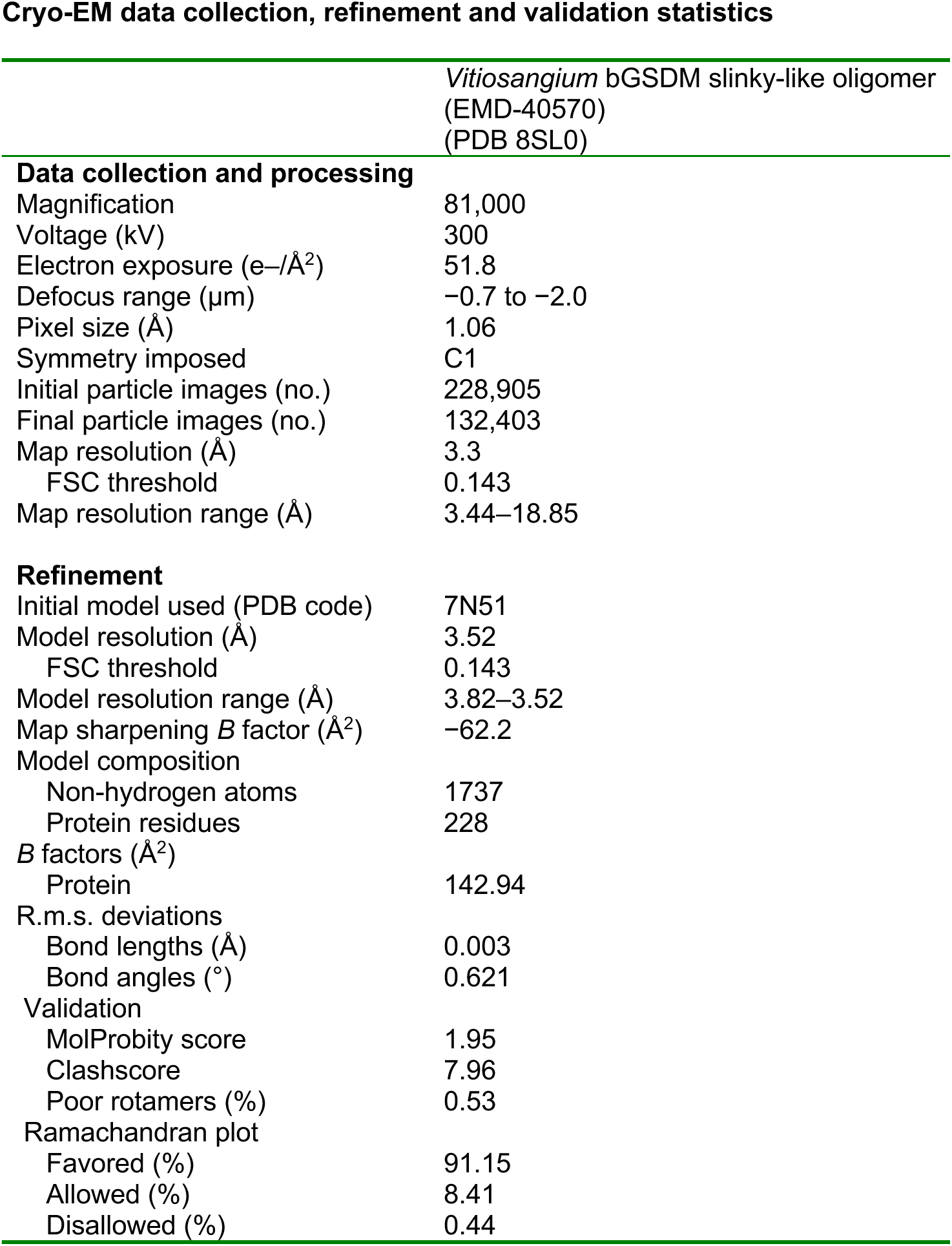
Cryogenic-electron microscopy data summary table. Table contains details of all cryo-EM data collection, processing, and refinement statistics used in this study.

**Supplementary Table 1.** Restraints used during energy minimization (EM) and stepwise equilibration (EQ) in kJ mol^−1^ nm^−2^.

| Step | Time [ns] | Backbone | Sidechains | Lipid headgroup (z) | Lipid dihedrals |
| --- | --- | --- | --- | --- | --- |
| EM |  | 4000 | 2000 | 1000 | 1000 |
| EQ 1 | 50 | 2000 | 1000 | 50 | 0 |
| EQ 2 | 50 | 500 | 100 | 0 | 0 |

**Supplementary Table 2.**
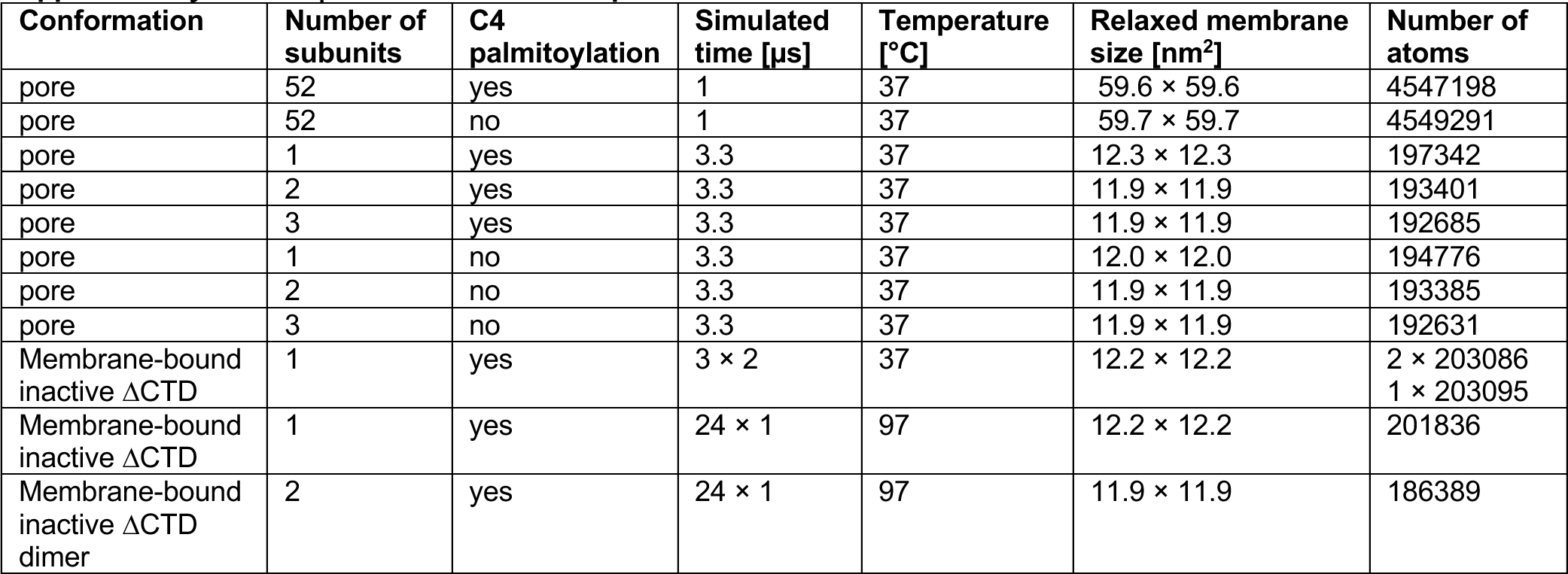
Overview of bGSDM production simulations.

